# Clonal and developmental relationships between Tph and Tfh cells in patients with SLE and in murine lupus

**DOI:** 10.1101/2025.01.27.635189

**Authors:** Takanori Sasaki, John Sowerby, Yinan Xiao, Runci Wang, Snigdha Samarpita, Tatsuki Abe, Kathryne E. Marks, Alice Horisberger, Yidan Gao, Ridwan B. Rashid, Pui Y. Lee, Yujie Qu, Marc A. Sze, Stephen E. Alves, Marc C. Levesque, Keishi Fujio, Karen H. Costenbader, Peter T. Sage, Deepak A. Rao

**Author notes:** equal contribution (Takanori Sasaki, John Sowerby). **Correspondence:** Deepak Rao, MD PhD, Division of Rheumatology, Inflammation, Immunity Brigham and Women’s Hospital, Hale Building for Transformative Medicine, 6002R 60 Fenwood Road, Boston, MA, USA.

## Abstract

Pathologic T cell-B cell interactions drive disease in systemic lupus erythematosus (SLE). The T cells that activate B cell responses include T peripheral helper (Tph) and T follicular helper (Tfh) cells, yet the developmental and clonal relationships between these B cell-helper T cell populations are unclear. Here we use T cell receptor (TCR) profiling to demonstrate substantial clonal overlap between Tph and Tfh cells in the circulation of patients with SLE. Expanded Tph and Tfh cell clones persist over the course of 1 year in patients with a new diagnosis of SLE, and clones are observed to shift both from Tfh to Tph cells and from Tph to Tfh cells over time. High resolution analysis of CXCR5^-^ PD-1^hi^ CD4 T cells, a common approach to identify Tph cells, sorted from blood of SLE patients revealed considerable heterogeneity among these cells and highlighted a subpopulation of cells with transcriptomic features characteristic of activated B cell-helper Tph and Tfh cells. This cell population, marked by expression of *TOX* and *CXCL13,* was found in both sorted Tph and Tfh cells, and was clonally linked in these two populations. Analysis of B cell-helper T cells in murine pristane-induced lupus also identified clonally linked yet transcriptionally distinct Tph and Tfh populations in spleen and lung. CXCR5 fate mapping demonstrated that Tph cells could arise through both CXCR5-experienced and CXCR5-independent differentiation routes. T cell-specific loss of Bcl6 prevented the accumulation of Tfh cells and reduced the accumulation of Tph cells, demonstrating a requirement for Bcl6 in the development of both populations. Together, these observations reveal close clonal and developmental relationships between Tph and Tfh cells while demonstrating that Tph cells can arise through multiple differentiation routes.

## Introduction

Systemic lupus erythematosus (SLE) is a systemic autoimmune disease that can cause immune-mediated injury to multiple organs (*1*). Pathologic interactions of T cells and B cells and production of autoantibodies are core features of SLE immunopathology (*2*). Production of autoantibodies typically requires both autoreactive B cells and help from autoreactive T cells, which express T cell receptors (TCR) capable of recognizing the targeted self-antigens. Analyses of circulating T cells from patients with SLE have demonstrated an increase in circulating CXCR5^+^ PD-1^hi^ T follicular helper (Tfh) cells, as well as a prominent increase in circulating CXCR5^-^ PD-1^hi^ T peripheral helper (Tph) cells (*3–7*). Both Tfh and Tph cells are considered "B cell-helper T cells," as they are capable of recruiting and activating B cells through production of the B cell chemoattractant CXCL13 and the cytokine IL-21, which is required to induce B cell differentiation into plasmablasts (PB) and age-associated B cells (ABC)(*3*, *8*, *9*). Expansion of circulating Tph and Tfh cells has been observed in multiple SLE patient cohorts, generally with higher frequencies in patients with active disease (*4–6*, *10*, *11*). Longitudinal studies have indicated that Tfh cell expansion in blood is most evident early in disease and may decrease over time, while Tph cell expansion persists in established disease (*12*). In a cohort of patients with lupus nephritis, the frequency of Tph cells, but not Tfh cells, correlated with both disease activity and the frequency of circulating ABCs (*4*). The developmental relationship between Tph and Tfh cells remains largely unknown.

One challenge in evaluating the relationships between these cell populations is that distinguishing B cell-helper T cells from other activated T cell populations phenotypically remains difficult. Tfh cells express CXCR5, a chemokine receptor that detects the ligand CXCL13 to induce cell migration into secondary lymphoid organ follicles (*13–16*). More refined populations of active Tfh cells or germinal center Tfh cells are defined by high levels of expression of PD-1, ICOS, BCL6, and CXCR5 (*14–17*). Tfh cells require the transcription factor BCL6, such that absence of BCL6 prevents accumulation of Tfh cells in vivo (*18–20*). Identifying Tph cells, which can be functionally defined as B cell-helper T cells that lack CXCR5 and instead express migratory programs targeting non-lymphoid tissues, is more challenging. Tph cells are often quantified in patient samples as CXCR5^-^ PD-1^hi^ CD4 T cells (*3–6*); however, this gating strategy is likely to capture heterogeneous populations that include other activated T cells. In patients with SLE and rheumatoid arthritis (RA), the CXCR5^-^ PD-1^hi^ T cell population is enriched for a B cell-helper transcriptomic signature, and these cells provide help to B cells in vitro, indicating a clear B cell-helper phenotype within this cell population in these diseases (*3*, *4*). However, the proportion of B cell-helper T cells versus other T cell effectors among circulating PD-1^hi^ T cells may vary across diseases. Single cell genomic studies have demonstrated transcriptomically or epigenetically separate populations of Tph and Tfh cells within inflamed tissue (*21*, *22*), both with B cell-helper transcriptional signatures, yet the ability to distinguish these cells transcriptomically in blood has not been clearly established.

In addition to transcriptomic parallels, developmental links between T cell populations in human studies can be inferred from TCR profiling, which provides a tool to track T cell clones across space, time, and phenotype. Each T cell produced in the thymus expresses a unique TCR, which is passed on to all progeny from that T cell clone. The TCRs can thus be utilized as molecular barcodes, enabling the tracking of each T cell’s developmental trajectory. This clonal tracking approach can also be applied to longitudinal samples, making it possible to trace how specific T cell subsets evolve over time and into different subsets. Applied to samples from autoimmune disease patients, TCR-based clonal tracking has been used to link immune responses across anatomical compartments, follow disease-associated clones over time, and infer relationships between distinct T cell populations. Studies in type I diabetes and Sjögren disease identified shared TCRs in target tissues and blood and expansion of antigen-specific TCRs during progression to disease (*23–25*). Longitudinal analysis demonstrated the persistence of expanded T cell clones in the circulation of patients with systemic sclerosis over time (*26*). More recently, a large-scale TCR repertoire analysis in SLE revealed increased clonotype sharing between effector Treg and Th1 cells, suggesting plasticity between disease-associated T cells (*27*).

Here, we report detailed profiling of TCRs of specific T cell populations from the circulation of patients with SLE, including Tph cells, Tfh cells, T regulatory cells (Tregs), and comparator T cell populations, obtained at diagnosis, 6 months, and 1 year. We combined this with single-cell RNA/TCR profiling for additional cellular resolution to interrogate clonal relationships between Tph and Tfh cell populations in SLE patients. We further profiled Tph and Tfh cells from lung and spleen of mice with pristane-induced lupus to track and manipulate the development of these cells in vivo. These studies demonstrate a strong TCR repertoire overlap between Tph and Tfh cells in SLE patients and in murine pristane-induced lupus across different time points and different tissues, with fate-mapping and interventional evidence further supporting a tight, though non-exclusive, developmental link between Tph and Tfh cells.

## Results

### Persistent shared clones in bulk sorted Tph and Tfh cells from patients with SLE

To compare the TCR repertoires of specific T cell populations and their changes over time in SLE, we performed TCR sequencing on T cell subsets sorted from PBMC of 9 patients with a new diagnosis of SLE. Samples were collected within 6 months of diagnosis, and then 6 months and 12 months later. TCR repertoire analysis was conducted on 7 memory CD4 T cell populations (CD25^+^ CD127^-^ [Treg], CXCR5^-^PD-1^hi^, CXCR5^+^ PD-1^hi^, CXCR5^-^ PD-1^int^, CXCR5^+^ PD-1^int^, CXCR5^-^ PD-1^low^, and CXCR5^+^ PD-1^low^) (Figure 1A). CXCR5^-^ PD-1^hi^ cells, which contain Tph cells, exhibited the highest degree of clonal expansion, with persistent clonal expansion at each timepoint (Figure 1B, Extended Data Figure 1A). Comparison of the overlap of TCR repertoires across different subsets demonstrated substantial clonal overlap across T cell subsets within each patient. We observed significant TCR repertoire overlap between CXCR5^-^ PD-1^hi^ and CXCR5^+^ PD-1^hi^ cells (Figure 1C, indicated as ‘a’), as well as between CXCR5^-^ PD-1^hi^ and CXCR5^-^ PD-1^int^ cells (indicated as ‘b’), between CXCR5^-^ PD-1^int^ cells and CXCR5^-^ PD-1^low^ cells (indicated as ‘c’), and between CXCR5^+^ PD-1^hi^ and CXCR5^+^ PD-1^int^ cells (indicated as ‘d’). The significant repertoire overlap between CXCR5^+^ PD-1^hi^ and CXCR5^-^ PD-1^hi^ populations, which are well separated cytometrically, supports a developmental relationship between Tph and Tfh cells. TCRs of Tregs showed little clonal overlap with any other T cell subset. TCR profiling of T cell subsets from a cohort of 10 non-inflammatory control patients demonstrated a similar pattern of TCR repertoire overlap across cell subsets, indicated that the clonal relationships between T cell populations are not disease-specific (Extended Data Figure 1B). There was minimal TCR overlap across different patients, as expected (Extended Data Figure 1C).

**Figure 1.**
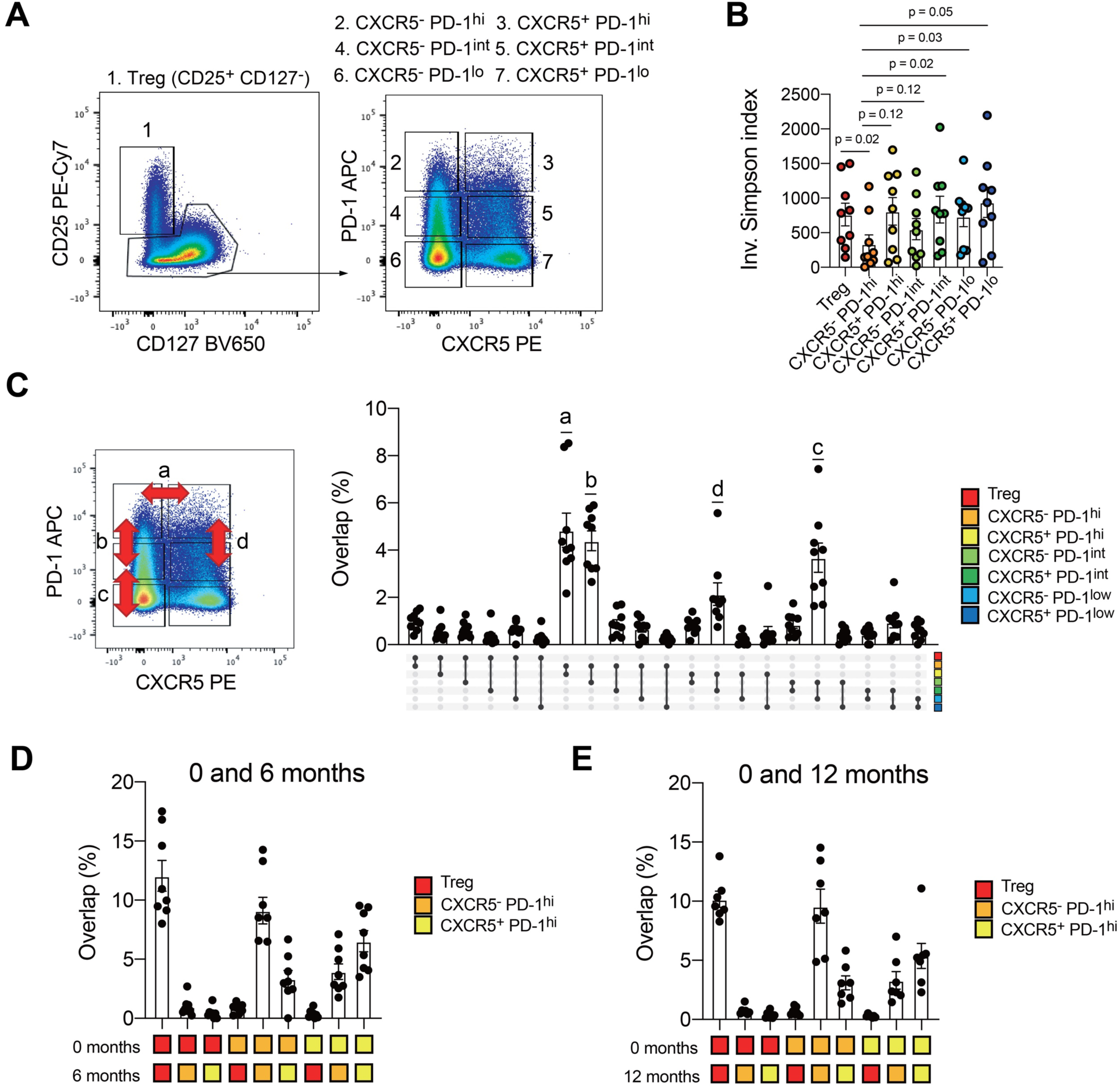
Longitudinal changes of TCR repertoire features in SLE. **A.** Schematic of TCR repertoire analysis in SLE. Bulk TCRs of Tph cells, Tfh cells, Tregs, and 4 other comparator CD4 T cell subsets were generated from blood of 9 patients with a new diagnosis of SLE, followed across 3 timepoints and 10 control donors. **B.** TCR clonality of T cell subsets (SLE: n = 9). TCR clonality was assessed by inverse Simpson Index (downsampled to 5000 reads). P-values by Wilcoxon test. **C.** TCR repertoire overlap between Tph cells, Tfh cells, Tregs, and 4 other comparator CD4 T cell subsets in SLE (n = 9, downsampled to 5000 reads). TCR overlap was calculated by the repOverlap function in the immunarch package with the “overlap” method. **D.** TCR repertoire overlap between Tph cells, Tfh cells, and Tregs across baseline and 6 months (SLE: n = 9, downsampled to 5000 reads). **E.** TCR repertoire overlap between Tph cells, Tfh cells, and Tregs across baseline and 1 year (SLE: n = 7, downsampled to 5000 reads).

Longitudinal analyses of T cell subsets from the SLE cohort indicated that CXCR5^-^ PD-1^hi^, CXCR5^+^ PD-1^hi^, and Treg populations have expanded TCR clones that persist over at least the first year after diagnosis (Figure 1D, E). The repertoire overlap between CXCR5^-^ PD-1^hi^ and CXCR5^+^ PD-1^hi^ cells was consistently observed when comparing samples from diagnosis versus 6 months later, and comparing diagnosis versus 1 year later (Figure 1D, E). In addition, unique TCR clones from CXCR5^+^ PD-1^hi^ cells, not shared with CXCR5^-^ PD-1^hi^ cells at baseline, were also detected in CXCR5^-^ PD-1^hi^ cells, and vice versa, at 6 months and 1 year (Extended Data Figure 1D), suggesting potential phenotype transitions within clones. Treg clones remained unique to the Treg subset without substantial appearance in CXCR5^-^ PD-1^hi^, CXCR5^+^ PD-1^hi^, or other T cell subset repertoires over time.

### A cluster of transcriptomically active B cell-helper T cells within PD-1^hi^ cell populations

To evaluate TCRs of these T cell populations at higher cellular resolution, we conducted scRNA/TCRseq on T cell subsets from 3 of the SLE patients in this cohort using PBMC collected at 18 months after enrollment. We selected 3 patients with a high degree of clonality in PD-1^hi^ cells for analysis. To unambiguously distinguish between CXCR5^-^ PD-1^hi^ cells (containing Tph) and CXCR5^+^ PD-1^hi^ cells (Tfh), we sorted CXCR5^-^ PD-1^hi^, CXCR5^+^ PD-1^hi^, CXCR5^-^ PD-1^int^, CXCR5^+^ PD-1^int^, and PD-1^low^ cells from non-Treg memory CD4 T cells, and then generated scRNA/TCR-seq data on each sorted population. UMAP visualization of all cells identified 9 clusters (Figure 2A-B): C0 (Tfh), C1 (Th1), C2 (Th2), C3 (cytotoxic T lymphocyte; CTL), C4 (Th17), and C5 (TOX^+^ CXCL13^+^). Over 50% of CXCR5^+^ PD-1^hi^ cells clustered in C0. In addition, 13% of CXCR5^+^ PD-1^hi^ cells clustered in C5 and 12% clustered in C8 (*GZMK*^+^). The C5 cluster expressed several genes associated with B cell-helper function, including *CXCL13*, *IL21*, *CD200*, and *ICOS*. In contrast to Tfh cells, CXCR5^-^ PD-1^hi^ cells were more heterogeneous, with cells localized to several different clusters. 16% of these cells localized to C5, such that only cells sorted as PD-1^hi^ Tph or Tfh cells substantially contributed to the C5 cluster representing active B cell-helper T cells. Consistent with previous reports (*10*, *28*), BCL6 gene expression was not detectable in blood CD4 T cells; however, inference of BCL6 transcription factor activity revealed significantly higher BCL6 activity in C0, supporting the annotation of C0 as a Tfh cluster (Figure 2C).

**Figure 2.**
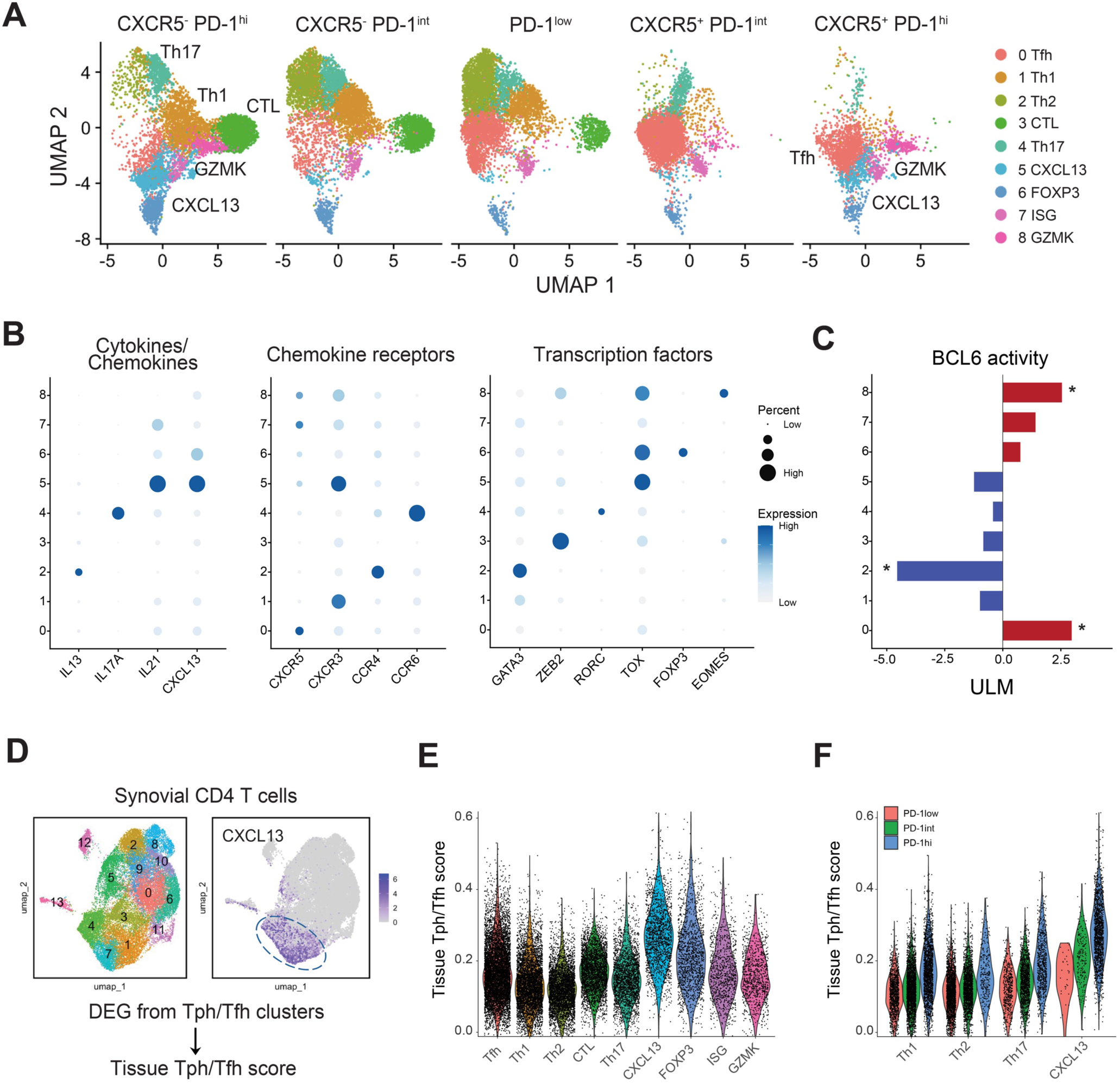
Single cell RNA-seq revealed activated B cell helper T cells. **A.** UMAPs showing cluster composition of scRNA-seq profiles of sorted CXCR5^-^ PD-1^hi^, CXCR5^+^ PD-1^hi^, CXCR5^-^ PD-1^int^, CXCR5^+^ PD-1^int^, and PD-1^low^ cells from blood CD4 T cells (n = 3 SLE donors). **B.** Gene expression profiles of each cluster in SLE blood CD4 T cell scRNA-seq data. **C.** BCL6 TF activity of each cluster. **D.** UMAP of RA synovial CD4 T cells and CXCL13, with scheme to define tissue Tph/Tfh gene signature score. **E.** Tissue Tph/Tfh gene signature score across SLE blood CD4 T cell clusters. **F.** Tissue Tph/Tfh score of Th1, Th2, Th17, and TOX^+^ CXCL13^+^ cells from sorted PD-1^low^, CXCR5^-^ PD-1^int^, and CXCR5^-^ PD-1^hi^ populations.

To confirm the identity of C5 cells as activated B cell-helper T cells, we established a tissue Tph/Tfh signature score based on the top 100 genes distinctively expressed by Tph/Tfh clusters from RA synovial tissue, where Tph cells were initially described (*3*, *29*) (Figure 2D). This Tph/Tfh signature score well highlighted the tissue Tph/Tfh clusters in kidney (*30*) and skin (*31*) in SLE (Extended Data Figure 1E). In addition, the Tph/Tfh signature score was highest in C5, indicating that this cluster expressed a prototype Tph/Tfh signature (Figure 2E). In addition, both PD-1^hi^ populations contributed to the C8 *GZMK*^+^ cluster. In contrast, a separate CTL cluster primarily contained cells from CXCR5^-^ PD-1^hi^ and CXCR5^-^ PD-1^int^ populations, without a contribution from PD-1^hi^ CXCR5^+^ cells. Across each of Th1, Th2, Th17, and TOX^hi^ CXCL13^hi^ clusters, CXCR5^-^ PD-1^hi^ tended to have higher scores for the Tph gene signature as compared to CXCR5^-^ PD-1^int^ or PD-1^low^ cells (Figure 2F).

### TCR clonal overlap between circulating Tph and Tfh cell populations

We then assessed the TCR clonal relationships across the finer T cell clusters revealed by scRNA-seq. C3 containing CTL exhibited the most clonal expansion (Figure 3A, B). The presence of these highly clonally expanded CTLs accounts for the high oligoclonality of CXCR5^-^ PD-1^hi^ cells observed in Figure 1.

**Figure 3.**
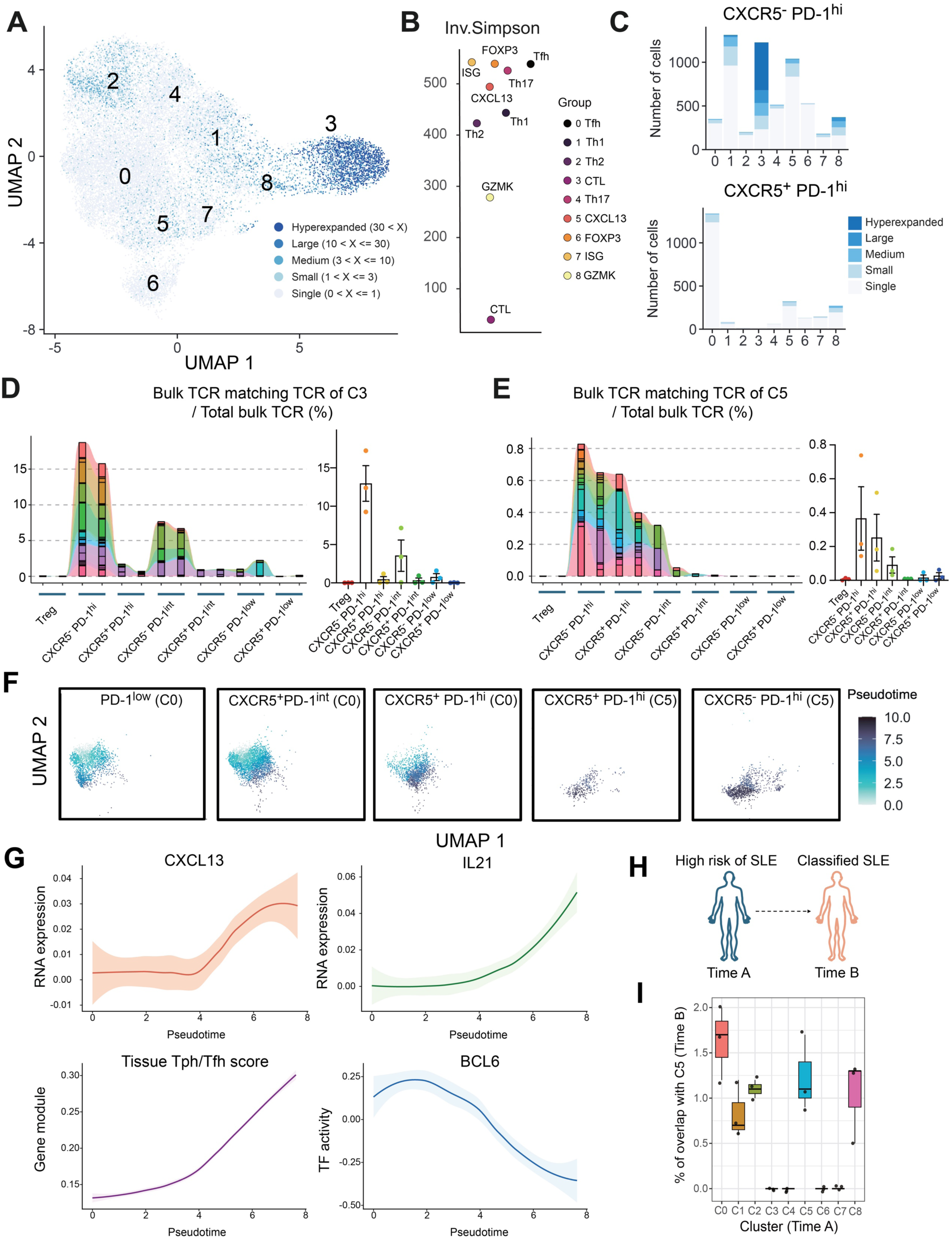
Clonal and developmental link between Tfh and Tph cells. **A.** Clonality of each cluster in SLE blood CD4 T cell scRNAseq data. Cells were classified based on the number of TCR clones detected. **B.** Inverse Simpson Index of each cluster. **C.** Clonality of each cluster within sorted CXCR5^-^ PD-1^hi^ and CXCR5^+^ PD-1^hi^ cell populations. **D.** Frequencies of TCR repertoire of C3 from patient SYL3 detected in bulk TCR-seq data at enrollment timepoint. Data from two technical duplicates are shown. **E.** Frequencies of TCR repertoire of C5 from SYL3 detected in bulk TCR-seq data at enrollment timepoint. **F**. Pseudotime analysis between C0 and C5 among sorted PD-1^low^, CXCR5^+^ PD-1^int^, CXCR5^+^ PD-1^hi^ and CXCR5^-^ PD-1^hi^ cells. **G**. Trend of gene expression, tissue Tph/Tfh signature score, and BCL6 regulon activity aligned along pseudotime. Colored lines represent LOESS fits across all cells, and shaded areas represent 95% confidence intervals for the fitted mean. **H**. Experimental scheme of the pre-SLE cohort. **I.** TCR overlap frequency between each subset at time A (pre-onset) and C5 at time B (post-onset).

Consistent with this, the extent of clonality of bulk CXCR5^-^ PD-1^hi^ pool was strongly correlated with the frequency of Tbet^+^ granzyme B^+^ CXCR5^-^ PD-1^hi^ CD4 T cells as quantified by mass cytometry analysis of the same samples (Extended Data Figure 1F). We then assessed the TCR clonality of CXCR5^-^ PD-1^hi^ or CXCR5^+^ PD-1^hi^ cells within each scRNA-seq cluster. C3 (CTL) from CXCR5^-^ PD-1^hi^ cells exhibited the most oligoclonality, while large, expanded clones could also be found in C5 (*TOX*^+^ *CXCL13*^+^), as well as C1 (Th1) and C8 (*GZMK*^+^) (Figure 3C).

To determine the specific cell clusters that account for the clonal overlap between CXCR5^-^ PD-1^hi^ and CXCR5^+^ PD-1^hi^ cells observed in the bulk TCR data, we used the scTCR data to determine the proportion of TCRs in each single cell cluster that was detected in bulk TCR-seq data from the enrollment time point. TCRs from cluster C3 (CTL) were predominantly associated with the bulk CXCR5^-^ PD-1^hi^ pool and were not found in the bulk CXCR5^+^ PD-1^hi^ pool (Figure 3D). In contrast, TCRs from C5 (TOX^+^ CXCL13^+^) were found in both bulk CXCR5^-^ PD-1^hi^ and CXCR5^+^ PD-1^hi^ pools (Figure 3E). This pattern of shared Tph/Tfh clones in C5 but not in C3 was consistent across the 3 SLE patients (Figure 3D, E, Extended Data Figure 1G), indicating that cells with a B cell-helper signature, and not CTL, account for the clonal overlap in TCRs of bulk CXCR5^-^ PD-1^hi^ and CXCR5^+^ PD-1^hi^ pools. In addition, TCRs from C1 (Th1) and C3 (CTL) were shared between the CXCR5^-^ PD-1^hi^ pool and the CXCR5^-^ PD-1^int^ pool (Extended Data Figure 2A). In total, these results indicate shared B cell-helper T cell clones between PD-1^hi^ Tph and Tfh cells, distinguishable from large CTL clones found exclusively in the CXCR5^-^ PD-1^hi^ pool and not in CXCR5^+^ PD-1^hi^ Tfh cells.

To interrogate the direction of differentiation between C0 and C5, we performed a pseudotime analysis. The pseudotime visualization indicated a trajectory from PD-1^low^ cells in C0 to CXCR5^+^ PD-1^int^ in C0, CXCR5^+^ PD-1^+^ in C0, CXCR5^+^ PD-1^+^ in C5, then CXCR5^-^ PD-1^+^ in C5, with gradually increased expression of the tissue Tph/Tfh signature, *CXCL13,* and *IL21,* but a decreased BCL6 transcription factor activity along this trajectory (Figure 3F, G). Taken together, these TCR overlap results and pseudotime analysis suggest a potential transition from Tfh to Tph cell phenotypes.

### Tph/Tfh cell clones are expanded prior to full disease onset

To evaluate whether clonal expansion of Tph and Tfh cells occurs prior to the full clinical manifestation of SLE, we evaluated CD4 T cells longitudinally from a set of patients who did not initially meet the ACR/EULAR 2019 classification criteria for SLE, but scored 4-9 points and were considered clinically at high risk of developing SLE. From this cohort, we identified 3 individuals who provided blood samples before (‘high risk of SLE’) and after meeting classification criteria for SLE (‘classified SLE’). We sorted CXCR5⁻ PD-1^hi^ and CXCR5⁺ PD-1^hi^ cells to generate scRNA/TCR-seq and projected the scRNA-seq data onto the reference SLE UMAP (Extended Data Figure 3A, B). Consistent with the SLE data (Figure 2A), the CXCR5⁺ PD-1^hi^ population from these patients was predominantly composed of C0 (Tfh), whereas sorted CXCR5⁻ PD-1^hi^ cells contained cells that clustered in C1 (Th1), C2 (Th2), and C4 (Th17). Again, the B cell helper population C5 was present in both sorted CXCR5⁺ and CXCR5⁻ PD-1^hi^ cell populations (Extended Data Figure 3C). TCR clonality analysis confirmed that the most clonally expanded populations were C3 (CTL) and C8 (GZMK) (Extended Data Figure 3D), though the proportion of CXCR5^-^ PD-1^hi^ cells that clustered in C3 was smaller in this dataset.

We then used the scTCR-seq data to determine whether clones expanded at the classified SLE timepoint were also present at the high-risk (pre-classified SLE) timepoint. Of 6310 expanded clones in sorted CXCR5⁻ PD-1^hi^ cells post-SLE classification, 61% were also detected at the high-risk, pre-classified timepoint. Similarly, of 2447 expanded clones in sorted CXCR5^+^ PD-1^hi^ cells post-disease classification, 55% were also detected at the high-risk, pre-classified timepoint. This indicates that expanded Tph and Tfh cell clones are present even before full manifestation of SLE disease. To further explore the potential origins of the clones in C5, we calculated the overlap of TCR sequences between post-onset C5 and each pre-onset cluster. Clonal overlap with post-onset C5 was observed across multiple pre-onset transcriptional states, including C0, C1, C2, C5, and C8, with the greatest overlap observed with C0 (Figure 3I). These findings indicate that clones occupying the C5 state after disease classification can be traced to multiple transcriptional states before disease onset, while highlighting C0 as a pre-onset state associated with these clones.

### TOX^+^ IL21^+^ CXCL13^+^ T cells represent an activated proliferative B cell-helper population in SLE

scRNA-seq analyses highlighted a population of T cells, captured in cluster C5, that expressed multiple B cell-helper features and was found specifically among CXCR5^-^ PD-1^hi^ and CXCR5^+^ PD-1^hi^ cells. We surmised that the scRNA-seq analysis revealed a population of functionally active B cell-helper T cells in the circulation that is otherwise difficult to visualize; therefore, we sought to identify specific markers to more precisely capture the C5 population by cytometry. Analysis of differentially expressed genes (DEG) revealed *TOX*, *CD200*, *ICOS*, and proliferation markers such as *MKI67* and *CDK14* as preferentially expressed in C5 (Figure 4A), consistent with an activated, proliferative B cell-helper population. We then evaluated expression patterns of these markers in a mass cytometry dataset generated from the same patient cohort (*12*). ICOS^+^ Ki67^+^ cells were found to be most abundant in gated CXCR5^-^ PD-1^hi^ cells and least abundant in PD-1^low^ cells, consistent with the scRNA-seq data (Extended Data Figure 2B, C, Figure 2A). Unsupervised clustering of cells in the mass cytometry dataset also identified a population of ICOS^+^ Ki67^+^ Tph cells (Figure 4B). To test their association with SLE, we applied a Covarying Neighborhood Analysis (CNA) (*32*), which demonstrated that ICOS^+^ Ki67^+^ Tph cells, Tregs, and granzyme K^+^ CD4 T cells were significantly enriched in SLE patients compared to controls (Figure 4B, Extended Data Figure 2D).

**Figure 4.**
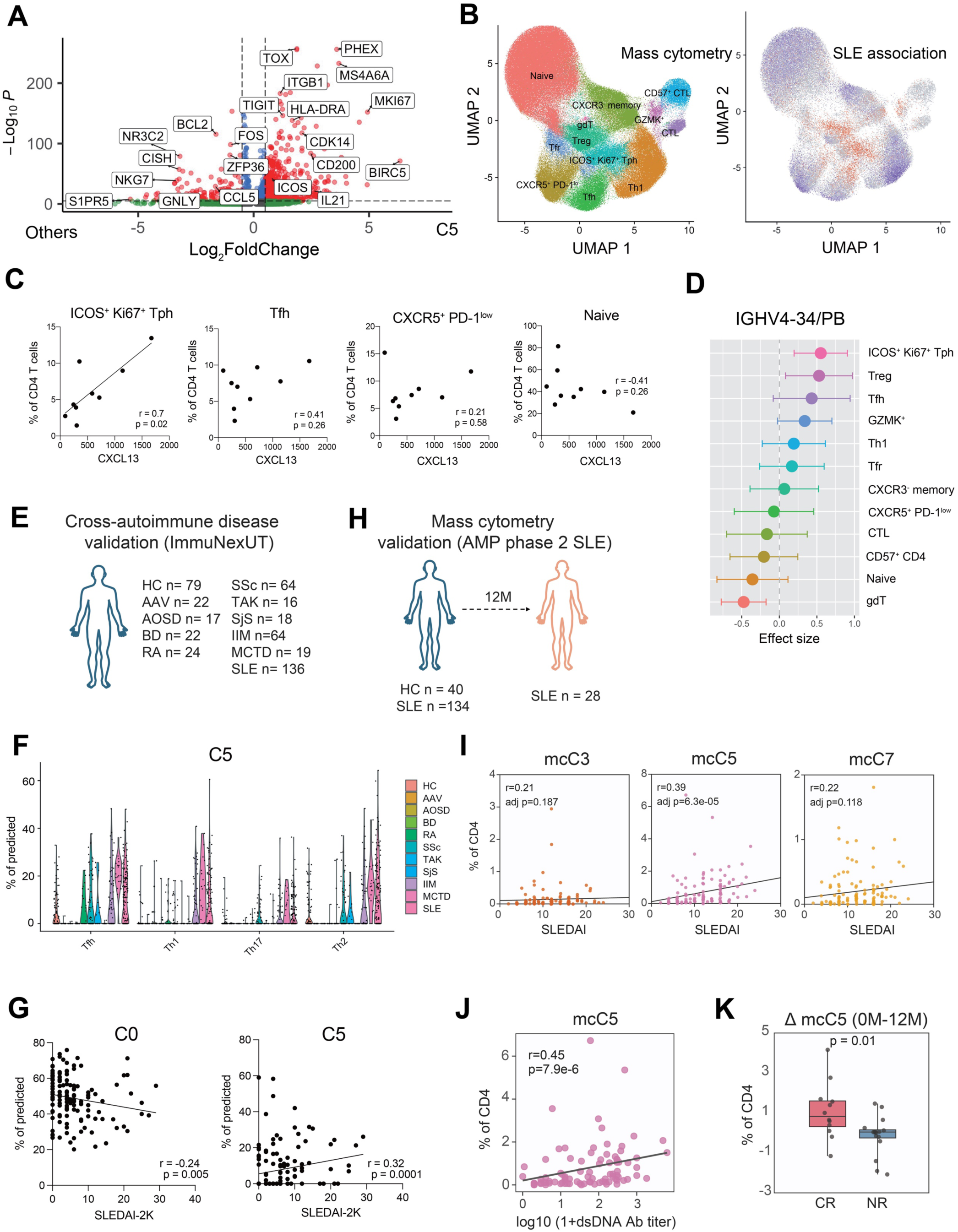
C5 is a strong B cell activator population. **A.** Volcano plot of DEGs between C5 and other clusters. Red: adj p values < 0.05, |log2FC| > 0.5, Green: adj p values > 0.05, Blue: adj p values < 0.05, |log2FC| < 0.5. **B. Left:** UMAP of blood CD4 T cells from mass cytometry of 14 non-inflammatory controls and 9 SLE patient samples. **Right:** CNA of mass cytometry data corrected for age and sex. Red indicates neighborhoods positively associated with SLE, and blue indicates negative association with SLE. **C.** Correlation between serum CXCL13 and the frequencies of ICOS^+^ Ki67^+^ Tph cluster, Tfh cluster, CXCR5^+^ PD-1^low^ cluster, and naive cluster at the baseline SLE samples. **D.** Linear mixed effect model analysis between mass cytometry clusters and the frequencies of IGHV4-34 on PB quantified by BCR-seq. 27 SLE patient samples (9 SLE patients x 3 time points) were used, with time as fixed effect. **E.** Overview of the ImmuNexUT bulk RNA-seq cohort used for deconvolution analysis (495 donors), including healthy controls (HC, n = 79) and patients with autoimmune diseases: ANCA-associated vasculitis (AAV, n = 22), adult-onset Still’s disease (AOSD, n = 17), Behçet’s disease (BD, n = 22), idiopathic inflammatory myositis (IIM, n = 64), rheumatoid arthritis (RA, n = 24), systemic sclerosis (SSc, n = 64), Takayasu arteritis (TAK, n = 16), mixed connective tissue disease (MCTD, n = 19), Sjögren syndrome (SjS, n = 18), and systemic lupus erythematosus (SLE, n = 136) (*35*, *36*). **F.** Estimated proportion of C5 cells among total cells, inferred by deconvolution analysis of ImmuNexUT bulk RNA-seq data. **G.** Spearman’s correlation of the estimated frequencies of C0 in Tfh and C5 in Th1 with SLEDAI-2K in ImmuNexUT study. (n = 136) (*36*). **H.** Mass cytometry validation cohort from 134 patients with SLE and 40 healthy controls (*7*), identifying Tph subclusters including C5. **I.** Correlation between the frequency of mcC3, mcC5, and mcC7 and SLEDAI-2K. **J.** Correlation between the frequency of mcC5, and anti-dsDNA antibody (n = 89). **K.** Longitudinal changes in the proportion of mcC5 cells in responders and non-responders following treatment in patients with lupus nephritis.

The proportion of ICOS^+^ Ki67^+^ Tph cells was also significantly associated with the serum CXCL13 level, whereas that of total CXCR5^+^ PD-1^hi/int^ Tfh cells was not (Figure 4C). ICOS^+^ Ki67^+^ Tph cells were also significantly associated with the frequency of IGHV4-34, a well-known autoreactive BCR-IGHV chain (*33*, *34*), quantified by bulk BCR-seq (Figure 4D). Together, these results indicate that tracking an ICOS^+^ Ki67^+^ Tph population captures an activated T cell population linked to overall T-B cell activation in SLE.

Next, to evaluate the C5 signature in additional cohorts of patients with SLE and other autoimmune diseases, we performed a deconvolution analysis using publicly available bulk RNA-seq data of CXCR3^+^ Th1, CCR4^+^ Th2, CCR6^+^ Th17, and CXCR5^+^ Tfh cells from 495 donors, including healthy controls and patients with 10 autoimmune diseases (Figure 4E) (*35*, *36*). Tph cells are not available in this dataset. C5 cells identified by deconvolution were observed in Th1, Th2, Th17, and Tfh populations, with the highest representation in the Tfh cell population. Compared to healthy controls, the proportion of C5 cells was increased in SLE, mixed connective tissue disease, and idiopathic inflammatory myositis patient samples (Figure 4F). Additionally, a transcriptomic dataset of 136 SLE patients (*35*, *36*) revealed that the frequency of C5 in Th1 was positively correlated with SLEDAI, whereas the frequency of C0 cells in Tfh cells was negatively correlated (Figure 4G). This inverse relationship between C0 and C5 suggests a potential differentiation shift from C0 to C5 in patients with higher disease activity.

For further validation, we analyzed an independent, large-scale mass cytometry dataset of PBMC from patients with lupus nephritis (*7*) (Figure 4H). Consistent with the single-cell transcriptomic analysis, substantial heterogeneity among CXCR5^-^ PD-1^hi^ CD4 T cells was also observed in the mass cytometry data (Extended Data Figure 4A). We attempted to quantify subclusters in the mass cytometry data of PD-1^hi^ CXCR5^-^ cells (designated mcC) that were analogous to the subclusters observed by scRNAseq (Fig 2A) using key markers in the mass cytometry panels that reflect features of the scRNA-seq subclusters, including Ki67 and ICOS (mcC5), FoxP3 and Helios (mcC6), GZMB (mcC3), GZMK (mcC8), CCR6 (mcC4), CXCR3 (mcC1), and MX1 (mcC7) (gated as in Extended Data Figure 4A). Among the CXCR5^-^ PD-1^hi^ subclusters, mcC1, mcC5, mcC6, mcC7, and mcC8 were significantly expanded in patients with SLE compared with healthy controls (Extended Data Figure 4B). The frequencies of mcC5 and mcC6 among CD4 T cells were positively correlated with SLEDAI, with the strongest correlation between mcC5 and disease activity (Figure 4I, Extended Data Figure 4C). mcC5 was also correlated with ds-DNA antibodies (Figure 4J). To determine whether these populations were associated with renal disease response to treatment, we also analyzed longitudinal samples obtained at baseline and 12 months after standard-of-care treatment. The frequency of mcC5 was significantly reduced in responders with lupus nephritis but remained unchanged in non-responders (Figure 4K). Together, these findings highlight a proliferating (Ki67^+^), ICOS^+^ PD-1^hi^ CXCR5^-^ Tph population as associated with disease activity across multiple, independent SLE cohorts.

### CXCR5^-^ PD-1^hi^ B cell-helper T cells accumulate in murine lupus

To evaluate whether similar heterogeneity and developmental links are observed among Tph cells in tissues, we evaluated PD-1^hi^ CD4 T cells in the murine model of pristane-induced lupus in C57BL/6 mice, a flexible model of lupus that involves activation of extrafollicular T-B cell interactions and can be induced in multiple strains (*37*). By two weeks after pristane treatment, a population of cells expressing high levels of PD-1 but lacking CXCR5, resembling human Tph cells, accumulated in spleen and peritoneal fluid (Figure 5A). At time points after two months, these cells became detectable in the lungs, peritoneal fluid, and lipogranulomas within the peritoneal cavity. This population was also present in the spleen, along with Tfh cells.

**Figure 5.**
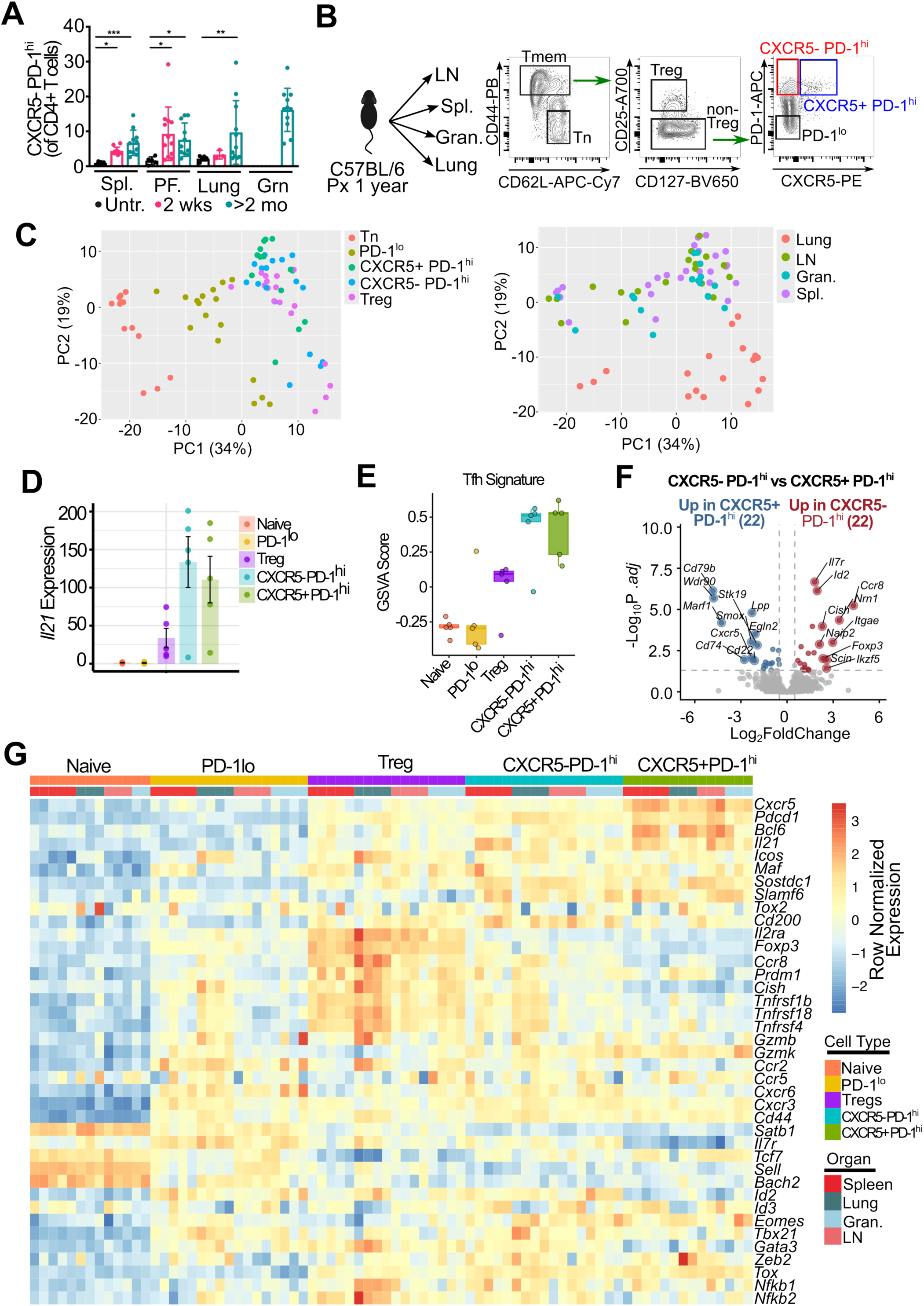
Spleen CXCR5^-^ PD-1^hi^ CD4 T cells in murine lupus have a B cell helper signature. **A**. Quantification of CXCR5^-^ PD-1^hi^ subset in pristane induced lupus in spleen (Spl), peritoneal fluid (PF), lung and granuloma (Grn) at indicated time-points. ANOVA results with indicated post-test significance. * p <0.05, ** p < 0.01, *** p <0.001, n= 3-4 per experiment. Pool of 3 independent experiments. **B**. Sorting schematic of the RNA-seq experiment. **C**. PCA analysis of sorted cell subsets grouped by cell type (left) or tissue (right). **D**. Splenic *Il21* expression of normalised counts across each of the sorted subsets. **E.** GSVA analysis using a Tfh gene signature across each of the different CD4 T cell subsets. **F**. DEG analysis of splenic CXCR5^-^ PD-1^hi^ vs CXCR5^+^ PD-1^hi^ subset. Some of the top differentially expressed genes are highlighted. **G**. Heatmap of gene expression for a custom gene list comprised of lineage, functional, migratory and activation markers. Normalised gene expression is plotted (row normalised).

To investigate the phenotype of these cells, we sorted T cell subsets from the spleen, draining lymph nodes, peritoneal granulomas, and lungs and performed RNA-seq (Figure 5B). Principal component analysis (PCA) revealed distinct clustering of cell types, with naïve cells, PD-1^low^ cells, and CXCR5^-^ PD-1^hi^/CXCR5^+^ PD-1^hi^/Treg cells forming separate clusters (Figure 5C). This clustering pattern was consistent across all tissues, although cells originating from the lungs clustered separately from other tissues. DEG analysis comparing CXCR5^-^ PD-1^hi^ cells from spleen with naïve T cells revealed upregulation of genes associated with B cell-helper function, including *Pdcd1*, *Icos*, and *Il21* in CXCR5^-^ PD-1^hi^ cells (Extended Data Figure 5A) with similar patterns observed in other tissues (Extended Data Figure 5B-C). Expression of *Il21* was equivalently elevated in the CXCR5^-^ PD-1^hi^ and CXCR5^+^ PD-1^hi^ T cell subsets (Figure 5D). GSVA analysis using a Tfh signature derived from known positive regulators of Tfh development (*38*) also showed equal enrichment between CXCR5^-^ PD-1^hi^ and CXCR5^+^ PD-1^hi^ T cell subsets (Figure 5E). Comparison of CXCR5^-^ PD-1^hi^ with CXCR5^+^ PD-1^hi^ cells revealed relatively few DEGs, indicating a phenotypic similarity between the two populations (Figure 5F, Extended Data Figure 6A). However, consistent with human Tph cells, murine CXCR5^-^ PD-1^hi^ cells exhibited a reduced ratio of *Bcl6* to *Prdm1* (Extended Data Figure 6B).

Heatmap analysis of z-scored expression of genes associated with B cell-helper T cells revealed that CXCR5^-^ PD-1^hi^ cells shared transcriptional features with CXCR5^+^ PD-1^hi^ cells, characterized by high expression of *Cxcr5*, *Pdcd1*, *Il21*, and *Icos* (Figure 5G). However, CXCR5^-^ PD-1^hi^ cells also expressed genes typically associated with other subsets, including *Prdm1*, *Gzmb*, and members of the Tnfrsf family, suggesting phenotypic heterogeneity within this population.

### Tph and Tfh cells in the spleen and lung are transcriptionally distinct but clonally related

To better resolve B cell-helper phenotypes within CXCR5^+^ PD-1^hi^ and CXCR5^-^ PD-1^hi^ CD4 T cell populations in this model, we analysed total CD4 T cells from the spleen and lung of mice 12 months after pristane treatment using scRNA/TCR-seq. UMAP visualization identified 12 CD4 T-cell clusters distributed across the spleen and lung (Figure 6A). One splenic cluster (cluster 2) and one lung cluster (cluster 4) showed high expression of genes associated with B cell-helper function, including *Il21*, *Cxcr5*, *Pdcd1*, *Bcl6* and *Cd200* (Figure 6B–C). Expression of *Il21* was relatively selective to these B cell-helper clusters (Extended Data Figure 7A). However, expression of individual B cell-helper genes varied within each cluster, suggesting within-cluster heterogeneity. We therefore re-clustered the splenic and lung B cell-helper populations separately. Cells within splenic cluster 2 segregated into four subpopulations (Figure 6D).

**Figure 6.**
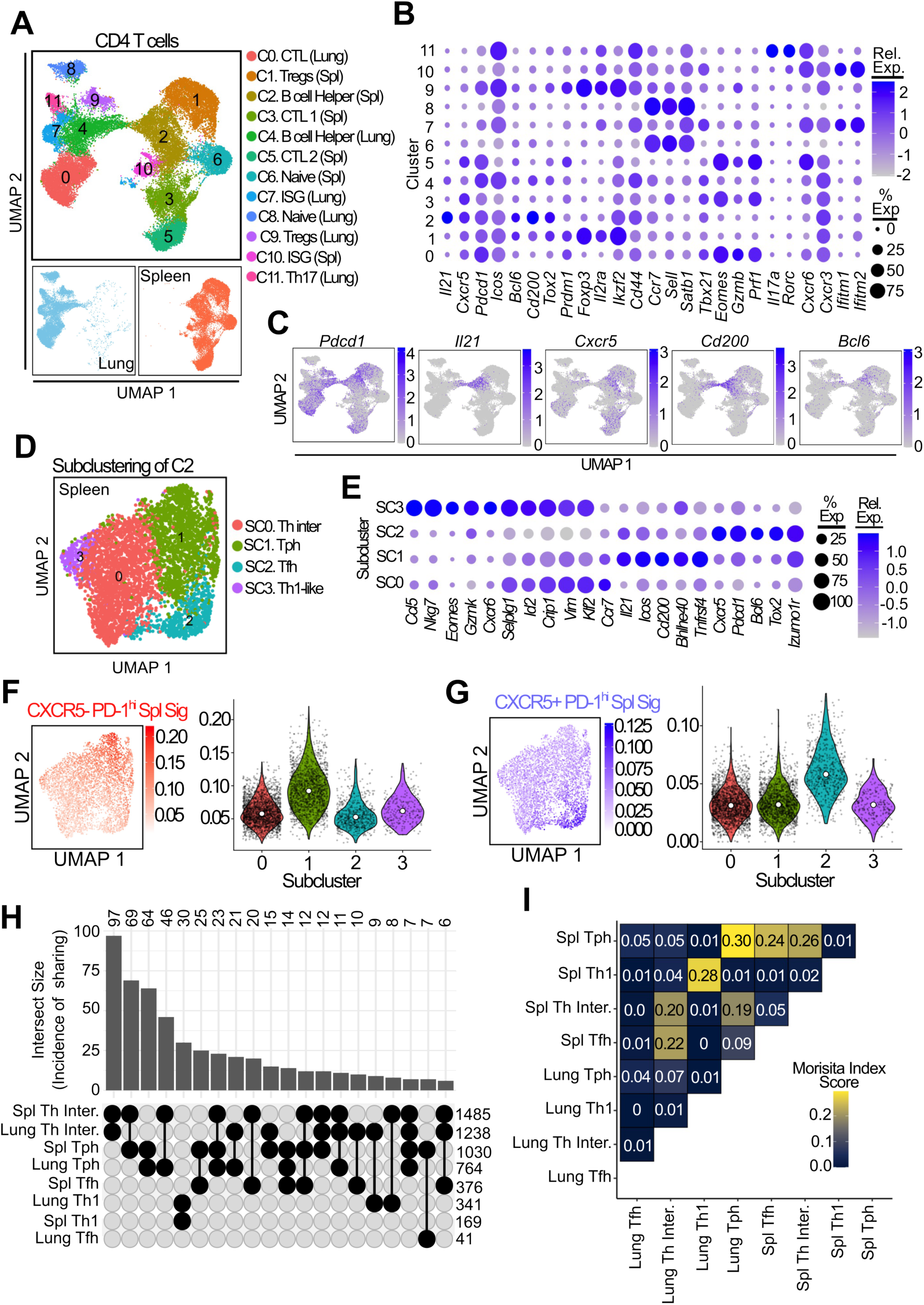
Single-cell RNA sequencing reveals phenotypic heterogeneity and shared clonotypes among B-cell-helper T-cell populations in murine lupus. **A.** UMAP visualization of integrated CD4 T cells from spleen and lung, with clusters annotated by cell state and tissue of origin. Lower panels show the distribution of lung- and spleen-derived cells. **B.** Dot plot showing expression of selected genes associated with naïve, regulatory, T helper, and B-cell-helper states across the identified clusters. Dot size represents the percentage of cells expressing each gene, and colour indicates scaled average expression. **C.** Feature plots showing expression of *Pdcd1, Il21, Cxcr5, Cd200,* and *Bcl6* across the integrated dataset. **D.** UMAP visualization of subclusters identified following reclustering of the splenic B-cell-helper cluster (C2), annotated as T helper intermediate (SC0), Tph (SC1), Tfh (SC2), and Th1-like (SC3) populations. **E.** Dot plot showing expression of selected genes across the four splenic B-cell-helper subclusters. **F, G.** UMAP and violin plots showing module scores across the splenic B-cell-helper subclusters using gene signatures derived from bulk-sorted CXCR5⁻PD-1^hi^ cells (**F**) or CXCR5⁺PD-1^hi^ cells (**G**). White points indicate the median score within each subcluster. **H.** UpSet plot showing the incidence and size of clonotype intersections among Tph, Tfh, Th1, and T-helper-intermediate populations from the spleen and lung. Horizontal values indicate the total number of clonotypes in each population, and vertical bars indicate the number shared by the population combinations shown below. **I.** Pairwise Morisita similarity indices quantifying clonotype repertoire overlap among the corresponding spleen and lung T-cell populations.

Subcluster 2 showed a clear Tfh phenotype, with the highest expression of *Bcl6*, *Tox2* and *Cxcr5* (Figure 6E). In contrast, subcluster 1 had the highest expression of *Il21, Icos, Cd200*, *Tnfrsf4* and the transcription factor *Bhlhe40*, demonstrating features shared with human Tph cells (*39*). Subcluster 0 showed an intermediate B cell-helper phenotype, whereas subcluster 3 exhibited a Th1-like state. Re-clustering of lung cluster 4 identified a similar organization, although the distinction between the Tph and intermediate T-helper populations was less pronounced than in the spleen (Extended Data Figure 7B–C).

To determine whether the transcriptional differences between Tph- and Tfh-like subclusters paralleled those observed between the corresponding bulk-sorted populations, we scored cells using gene signatures derived from bulk sorted CXCR5^-^ PD-1^hi^ and CXCR5⁺ PD-1^hi^ CD4 T cells from spleen of pristane-treated mice (Figure 5B, Supplementary Table 1). The CXCR5^-^ PD-1^hi^ signature was most highly enriched in splenic subcluster 1, whereas the CXCR5⁺ PD-1^hi^ signature was highest in subcluster 2, supporting their identification as Tph and Tfh cells, respectively (Figure 6F–G). A similar pattern was observed in the lung: the CXCR5^-^ PD-1^hi^ signature was enriched in the Tph subcluster, whereas the CXCR5⁺ PD-1^hi^ signature was highest in the Tfh subcluster (Extended Data Figure 7D–E).

We next assessed the correspondence of these populations with independently defined human Tph and Tfh states. A human Tph signature derived from RA synovial Tph cells showed higher expression in the murine Tph clusters in both spleen and lung, while a human Tfh signature derived from tonsil Tfh cells showed higher expression in the murine Tfh clusters (Extended Data Figure 7F, Supplementary Table 2). GSVA of per-mouse, per-subcluster pseudobulk profiles further showed that the signature of the human blood Tph population (C5) identified earlier in this study (Supplementary Table 3) was most enriched in the mouse Tph subclusters in both spleen and lung (Extended Data Figure 7G). Direct differential-expression analysis of spleen Tph and Tfh cells identified higher expression of Tfh-associated genes, including *Cxcr5*, *Tox2, Bcl6, Pou2af1* and *Ascl2*, in the Tfh population. Conversely, Tph cells showed increased expression of *Ahnak, Anxa6, Id2, Bhlhe40,* and *Tnfrsf4* (Extended Data Figure 7H). Consistent with these findings, GSEA demonstrated significant enrichment of a human Tph signature among genes upregulated in murine Tph cells and enrichment of a human tonsillar Tfh signature among genes upregulated in murine Tfh cells (Extended Data Figure 7I). Together, these analyses identify transcriptionally separable Tph and Tfh populations within the B-cell-helper compartments of both the spleen and lung.

We then used the paired scTCR-seq data to examine the clonal relationships between these transcriptionally defined populations. Clonally expanded cells were distributed across multiple CD4 T cell states, with the greatest expansion observed within cytotoxic T cell populations (Extended Data Figure 8A). Extensive TCR sharing was observed among splenic and lung B cell-helper populations: clonotypes from both splenic Tph and Tfh cells were detected in the lung B-cell-helper compartment, while lung Tph and Tfh clonotypes were shared most frequently with splenic B cell-helper cells, although relatively few lung Tfh cells were recovered (Figure 6H, Extended Data Figure 8B–E). Morisita overlap index identified the strongest cross-tissue repertoire overlap between splenic and lung Tph cells and also demonstrated substantial overlap of splenic Tph cells with splenic Tfh and intermediate B-cell-helper populations (Figure 6I). In contrast, splenic and lung Th1 populations showed a distinct pattern of repertoire overlap, consistent with a clonal structure separate from the Tph–Tfh compartment. Together, these findings demonstrate clonal relationships among Tph, Tfh and related B-cell-helper states within and across tissues, consistent with shared developmental histories and/or transitions between these populations.

### CXCR5 lineage tracing reveals distinct routes to a Tph phenotype

To determine if Tfh cells can acquire a Tph-like phenotype, we first sorted CXCR5⁺ PD-1^hi^ Tfh cells and stimulated them in vitro for 5 days. Following stimulation, ∼30% of the cells lost CXCR5 while retaining high PD-1 expression, indicating that Tfh cells can transition to a CXCR5⁻ PD-1^hi^ population resembling Tph cells (Extended Data Figure 9A).

To track the potential transition of CXCR5^+^ T cells into a CXCR5^-^ Tph phenotype in vivo, we generated *Cxcr5-Cre Rosa26-LSL-YFP* lineage-reporter mice. In this model, cells that express Cxcr5 undergo permanent YFP labelling, allowing cells with current or prior CXCR5 expression to be identified even if they subsequently lose surface CXCR5 (Extended Data Figure 9B). Control experiments demonstrated that the reporter sensitively and specifically identified CXCR5-expressing cells, as ∼80% of B cells were YFP^+^, and that YFP labelling was not induced simply by T cell activation, as approximately half of PD-1^+^ CD69^+^ cells in pristane-treated mice were YFP^-^ (Extended Data Figure 9C and 9D). The model therefore enabled us to distinguish CXCR5⁻ PD-1^hi^ cells with a detectable history of CXCR5 expression from those without detectable lineage labelling.

We first quantified YFP labelling among splenic CD4 T cell populations defined by CXCR5 and PD-1 expression. Three months after pristane treatment, ∼90% of CXCR5^+^ PD-1^hi^ cells were YFP⁺, supporting the sensitivity of the reporter (Figure 7A). YFP was also detected in approximately 45% of CXCR5⁻ PD-1^hi^ cells, consistent with a proportion of these cells having previously expressed Cxcr5. In contrast, YFP was rare among CXCR5⁻ PD-1⁻ cells (Figure 7A).

**Figure 7.**
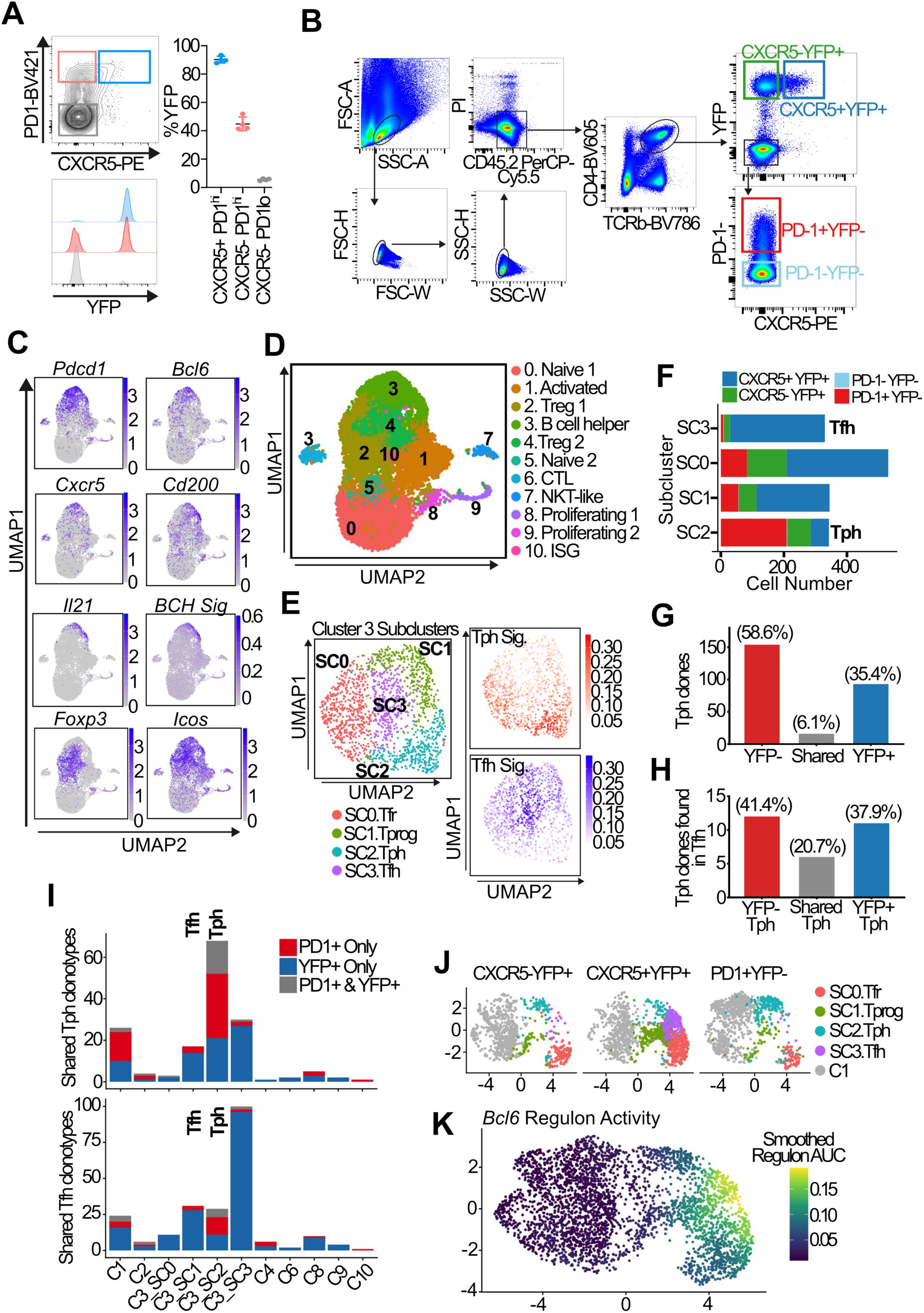
CXCR5 lineage tracing identifies distinct Tph and Tfh differentiation pathways in murine lupus model. **A.** Representative flow-cytometric analysis and quantification of YFP labelling among splenic CXCR5⁺PD-1^hi^, CXCR5⁻PD-1^hi^ and CXCR5⁻PD-1⁻ CD4 T cells. Points represent individual mice; horizontal lines and error bars show mean and SEM (n=4 mice). **B.** Flow-cytometric gating strategy used to sort CXCR5⁺YFP⁺, CXCR5⁻YFP⁺, PD-1⁺YFP⁻ and PD-1⁻YFP⁻ splenic CD4 T cells for scRNA/TCR-seq. **C.** Feature plots showing expression of *Pdcd1, Bcl6, Cxcr5, Cd200, Il21, Foxp3* and *Icos*, and a B-cell-helper signature score, across splenic CD4 T cells. **D.** UMAP of splenic CD4 T-cell clusters, including the B-cell-helper-enriched cluster 3. **E.** Reclustering of cluster 3 into Tfr (SC0), Tprog (SC1), Tph (SC2) and Tfh (SC3) states, with Tph and Tfh signature scores generated from 12mo scRNA-seq data shown alongside. **F.** Numbers of cells from each sorted population in the four cluster 3 subclusters. **G.** Numbers of Tph clonotypes detected only in YFP⁻ cells, in both YFP⁻ and YFP⁺ cells, or only in YFP⁺ cells. **H.** The same classification for clonotypes shared between the Tph and Tfh subclusters, according to their detection in YFP⁻ and YFP⁺ Tph cells. **I.** Numbers of Tph-associated (upper) and Tfh-associated (lower) clonotypes shared within other splenic CD4 T-cell clusters and subclusters. Colours indicate clonotypes detected only in PD-1⁺YFP-cells, only in YFP⁺ cells, or in both populations. **J.** UMAP of cluster 1 and cluster 3 subclusters, displayed separately for the CXCR5⁻YFP⁺, CXCR5⁺YFP⁺ and PD-1⁺YFP⁻ sorted populations. **K.** Smoothed SCENIC *Bcl6* regulon activity across the combined cluster 1 and cluster 3 UMAP.

To investigate the transcriptional relationships among these populations, we sorted CXCR5⁺YFP⁺, CXCR5⁻YFP⁺, PD-1⁺YFP⁻ and PD-1⁻YFP⁻ splenic CD4 T cells for scRNA/TCR-seq (Figure 7B). Expression of *Pdcd1, Bcl6, Cxcr5, Cd200, Il21* and *Icos*, together with a B cell-helper gene signature (Supplementary Table 4), identified a B cell-helper-enriched population within cluster 3 (Figure 7C–D). Reclustering of cluster 3 resolved four transcriptional states, annotated as Tfr, Tprog, Tph and Tfh (Figure 7E), similar to prior analyses of B6 mice (Figure 6). Tph- and Tfh-associated module scores (Supplementary Table 5) were again differentially enriched across the corresponding subclusters, supporting their identification as distinguishable B cell-helper states.

The Tfh subcluster was composed predominantly of cells sorted to be CXCR5⁺YFP⁺, consistent with current and historical CXCR5 expression. In contrast, the Tph subcluster contained a substantial contribution from CXCR5⁻YFP⁺ cells, indicating that cells that previously expressed CXCR5 can acquire a Tph transcriptomic identity (Figure 7F, Extended Data Figure 9E). In addition, the largest contribution to the Tph subcluster came from PD-1⁺ YFP^-^ cells, which have not previously expressed CXCR5. Thus, transcriptomically defined Tph cells can differentiate via both CXCR5+ and CXCR5-independent paths.

Evaluation of the associated TCR data demonstrated that among 263 clonotypes detected in the Tph subcluster, 154 were found only in YFP⁻ cells, 93 only in YFP⁺ cells and 16 in both (Figure 7G). 29 of the Tph clonotypes were also found in the Tfh subcluster, with a comparable number found in YFP^+^ and YFP^-^ derived Tph cells (Figure 7H).

Clonotypes associated with the Tph and Tfh states were also detected most prominently in activated cluster 1 (Figure 7I), identifying it as a candidate upstream population linked to both helper states. We therefore examined the transcriptional relationship between activated cluster 1 and the cluster 3 B-cell-helper states in a combined analysis. PD-1⁺YFP⁻ Tph cells occupied a region adjacent to cluster 1, while the pre-Tfh and Tfh states occupied a separate region of transcriptional space that connected to a different part of cluster 1 (Figure 7J). Together with the clonal relationships, this organization is consistent with distinct routes from an activated precursor population toward Tph and Tfh states. Finally, we examined Bcl6 regulon activity using SCENIC across this transcriptional landscape. Given the established role of Bcl6 in Tfh differentiation, we asked whether its activity also characterized the region occupied by PD-1⁺YFP⁻ Tph cells. Bcl6 activity was most apparent in the Tprog/Tfh region and was comparatively low along the cluster 1–Tph region occupied by PD-1⁺YFP⁻ cells (Figure 7K). Therefore, we next tested whether Tph and Tfh cells differed in their functional dependence on Bcl6.

### Bcl6 is required for the accumulation of both Tph and Tfh cells

To directly test the requirement for Bcl6 in the accumulation of Tph cells in pristane-induced lupus, we conditionally deleted Bcl6 from T cells using pristane-treated *Bcl6*^fl/fl^ CD4Cre⁺ B6 mice. CD4 T cells isolated from the spleen and lung three months after pristane treatment were analysed by scRNA/TCR-seq (Figure 8A).

**Figure 8.**
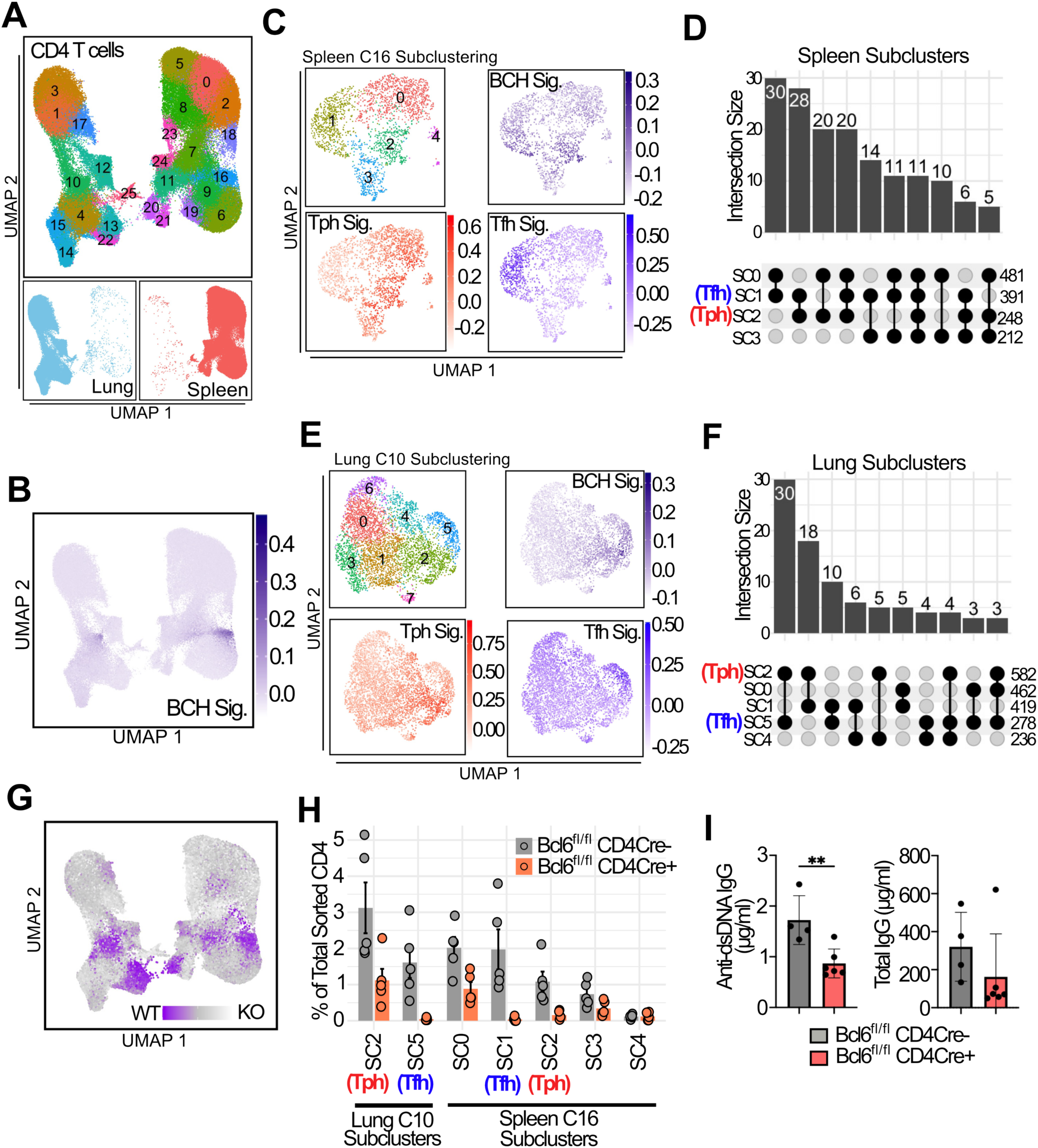
Absence of Bcl6 reduces accumulation of both Tph and Tfh cells in pristane-induced lupus. **A**. UMAP visualization of integrated splenic and lung CD4 T cells from the 3-month treatment cohort of *Bcl6*^fl/fl^ CD4Cre⁻ control and *Bcl6*^fl/fl^ CD4Cre⁺ conditional-knockout mice. Lower panels show the distribution of lung- and spleen-derived cells. **B**. B-cell-helper (BCH) module scores across the integrated dataset. The BCH signature was derived from genes upregulated in splenic cluster 2 of the 12-month scRNA-seq dataset. **C**. Reclustering of splenic cluster 16. UMAPs show the identified subclusters (top left), BCH module scores (top right), Tph-specific module scores (bottom left), and Tfh-specific module scores (bottom right). **D**. UpSet plot showing TCR clonotype sharing among the splenic cluster 16 subclusters. Horizontal values indicate the total number of clonotypes in each subcluster, and vertical bars indicate the numbers shared by the subcluster combinations shown below. **E, F**. Corresponding subclustering, module-scoring, and TCR-sharing analyses of lung cluster 10, (as described in **C** and **D)**. **G**. Covarying neighborhood analysis (CNA) showing the relative representation of *Bcl6*^fl/fl^ CD4Cre⁻ control and *Bcl6*^fl/fl^ CD4Cre⁺ cells across the integrated UMAP. Areas of genotype overrepresentation are colour-coded according to the inset scale. **H**. Proportion of control and Bcl6-deficient cells within the splenic cluster 16 and lung cluster 10 subclusters containing Tph- and Tfh-associated populations. **I**. Serum anti-dsDNA IgG and total IgG concentrations in *Bcl6*^fl/fl^ CD4Cre⁻ control and *Bcl6*^fl/fl^ CD4Cre⁺ mice, measured by ELISA. Points represent individual mice; n = 4–6 mice per group. Statistical significance was assessed using a two-sided Mann– Whitney U test; P < 0.01.

Using a similar analysis to prior experiments, we identified cell clusters with high expression of both a B cell-helper signature and high expression of *Il21 (*spleen cluster 16, lung cluster 10) (Figure 8B, Extended Data Figure 10A–B) and subclustered these populations, again identifying subclusters selectively enriched in Tfh (spleen subcluster 1, lung subcluster 5) or Tph signatures (spleen subcluster 2 and 3; lung subcluster 2). TCR clonal analyses again showed strong TCR overlap between spleen and lung Tph and Tfh clusters, similar to prior analyses (Figure 8D and 8F, Extended Data Figure 10C).

We used CNA to identify changes in cell population abundance following T cell-specific deletion of Bcl6. This analysis revealed a prominent depletion of B cell-helper T cells in both spleen (spleen cluster 16) and lung (lung cluster 10) in Bcl6-deficient mice compared with Bcl6^fl/fl^ CD4Cre⁻ controls (Figure 8G). Direct quantification of the relevant subclusters confirmed marked reductions in both Tph and Tfh cells in Bcl6-deficient mice (Figure 8H). These findings demonstrate that Bcl6 is required for the accumulation of both Tph and Tfh cells, despite the distinct CXCR5-associated histories and differentiation routes identified by lineage tracing.

Consistent with the reduction in B cell-helper T-cell populations, T cell-specific deletion of Bcl6 significantly reduced serum anti-dsDNA IgG concentrations (Figure 8I). Total IgG concentrations were also lower in Bcl6-deficient mice, although this difference was not statistically significant. Together, these findings identify Bcl6 as a shared requirement for Tph- and Tfh-cell development and suggest that the loss of these populations impairs pathogenic autoantibody production in pristane-induced lupus.

## Discussion

We performed detailed analyses of Tph and Tfh cells from the circulation of patients with early SLE profiled longitudinally and from pristane-induced murine lupus, in both cases demonstrating the shared transcriptomic and developmental features of these cell populations and their dynamics over time. Tph and Tfh cells from SLE patients show substantial overlap in their TCRs at multiple timepoints, consistent with past analyses of similarly sorted T cells from patients with HIV (*40*) and type-1 diabetes (*41*). Our scRNA-seq analyses of SLE T cell populations highlighted a specific cluster of PD-1^hi^ T cells with transcriptomic features of activation, proliferation, and B cell-helper function that likely reflects activated Tph and Tfh cells in circulation. These cells show clonal overlap, and trajectory and longitudinal clonal analyses suggest transitions between Tfh-related and Tph states, while also indicating that Tph cells may emerge from multiple transcriptional states. Expanded clones of both Tph and Tfh cells could be detected at multiple timepoints over 1 year, suggesting that disease-associated Tph and Tfh cell clones persist systemically in SLE patients. This persistence of clonally expanded Tph and Tfh cells may impose barriers to induction of stable tolerance by either traditional immunosuppressive agents or by B cell depletion.

Tph cells, as originally described in RA synovial tissue, expressed transcriptomic, cytometric, and functional features similar to Tfh cells, with a capacity to stimulate B cell differentiation in an IL-21-dependent manner (*3*). Cells with similar transcriptomic, cytometric, and functional features to RA synovial T cells can be found in the circulation as well (*42*), and many studies have reported quantification of circulating Tph and Tfh cells in autoimmune disease patient cohorts (*43*); however, identifying Tph cells in the circulation has been hampered by the difficulty in discriminating Tph cells from other activated T cell populations (*43*). Here, the recognition via scRNA-seq of a specific cluster of T cells with transcriptional features of activation, present in both sorted Tph and Tfh cell populations and showing shared clonality, provides a valuable tool in identifying circulating B cell-helper T cells and distinguishing these cells from PD-1^+^ cytotoxic T cells.

The human C5 cluster exhibited the highest expression of a Tph/Tfh signature score derived from tissue Tph/Tfh cells, along with high expression of *TOX*, *MKI67*, *ICOS*, *CXCL13*, and *IL21,* which strongly resembles Tph cells in the RA joints (*3*). Cluster C5 cell abundance correlated with both IGHV4-34^+^ plasmablasts and Ki67^+^ plasmablasts in SLE patients, suggesting that these T cells likely provide help to B cells. These findings align with previous observations that ICOS expression on PD-1^hi^ T cells enriches for B cell-helper function (*44*). C5 cluster cells also express TOX, a transcription factor that promotes exhaustion and PD-1 expression (*45*, *46*). Upregulation of TOX may suggest chronic antigen stimulation in these cells.

Parallel scRNA-seq analyses of CD4 T cells from the pristane-induced murine lupus model demonstrated heterogeneous populations of T cells expressing features associated with B cell help. These included conventional Tfh cells and transcriptionally distinct cells resembling human Tph cells, with substantial TCR overlap between the populations in both the spleen and lung. The range of B cell-helper T cell states observed in this model is consistent with previous reports describing populations that differ from prototypical Bcl6^hi^ CXCR5^hi^ Tfh cells. These include CXCR4⁺ extrafollicular helper T cells in murine lupus (*47*), CXCR5⁻ Tfh-like cells in the lungs of mice with chronic antigen-driven inflammation (*48*), and tissue-resident helper T cells identified in the lungs following influenza infection (*49*, *50*).

The substantial TCR overlap between Tph and Tfh cells demonstrates a close clonal relationship but does not establish the direction of differentiation or imply that all Tph cells arise from mature Tfh cells. The CXCR5 lineage-reporter model identified a population of CXCR5⁻ PD-1^hi^ Tph cells with permanent YFP labelling, providing evidence that some Tph cells develop after expressing CXCR5, potentially through a Tfh-precursor or Tfh state. This route may account for part of the clonal overlap between Tph and Tfh cells. It is also consistent with the persistence of circulating CXCR5⁻ PD-1^hi^ Tph cells as CXCR5⁺ PD-1^hi^ Tfh cells declined in patients with newly diagnosed SLE (*12*), and with the TCR repertoire overlap previously observed between Tph and Tfh cells in the NOD.AireGW/neuropathy model (*51*). Thus, in at least some settings, transition from a CXCR5-expressing state likely contributes to Tph cell accumulation.

However, approximately half of the transcriptionally defined Tph cells lacked detectable CXCR5 lineage labelling. Recognition of these cells supports a distinct route to the Tph state that does not require a detectable CXCR5-expressing history. This estimate must be interpreted in light of the sensitivity of the fate-mapping system; it is possible that T cells with weak or transient expression of CXCR5 may not produce sufficient Cre activity for recombination, which would lead to underestimation of the contribution of CXCR5-expressing cells to the Tph cell pool. Conversely, the high frequency of YFP labelling among CXCR5-expressing cells, appropriate labelling in genetic controls, and the absence of generalized YFP induction among activated CD4 T cells support the specificity of the reporter. In this context, the results support parallel routes to Tph differentiation, with contributions from both CXCR5+ and CXCR5-independent paths.

We find that in pristane-induced lupus, Bcl6 is required for Tph-cell accumulation. Although Bcl6 regulon activity was most apparent in the Tprog/Tfh region and comparatively low in the PD-1⁺YFP⁻ Tph region, T-cell-specific deletion of Bcl6 substantially reduced both Tph and Tfh cells in the spleen and lung of pristane-treated mice. Bcl6 can be induced early during CD4 T-cell activation, before establishment of a stable Tfh phenotype (*52*) and is also required for extrafollicular T-dependent antibody responses (*53*), raising the possibility that it acts at an earlier stage of Tph differentiation. Bcl6 expression has been reported to emerge during the transition from a Tfh progenitor state to a transitory state, before acquisition of the full effector Tfh program (*38*). One possibility is therefore that Tph and Tfh differentiation diverges during progression out of a progenitor-like state, after initiation of a Bcl6-associated program but before full effector Tfh differentiation. This may help explain the shared requirement for Bcl6 despite comparatively low Bcl6 regulon activity in the PD-1⁺YFP⁻ Tph region. Our experiments cannot directly determine whether the CXCR5-lineage-negative route itself requires Bcl6, because lineage history and Bcl6 dependence were assessed in separate models. Nevertheless, the marked overall depletion of Tph cells following Bcl6 deletion is consistent with a requirement extending beyond the CXCR5-lineage-positive population. Further studies will be required to define when Bcl6 acts along each pathway and how their relative contributions vary among disease settings.

This study is limited by the relatively small cohort of patients with early SLE analysed longitudinally and the inability to study tissue samples from patients. An additional limitation is that T follicular regulatory (Tfr) cells, which may have an important role in restraining autoreactive follicular immune responses, were not comprehensively characterized in this study. Future studies will be required to determine how Tfr populations interact with Tph and Tfh cells and influence pathogenic B-cell responses in SLE. Moreover, pristane-induced lupus is one of multiple lupus models that could be studied. In both the human and murine studies, the incomplete sampling of TCRs, assessing a small portion of the total T cells in each individual, may under-represent the extent of clonal sharing across T cell populations and timepoints.

In summary, these studies provide a delineation of the clonal dynamics of Tph and Tfh cells in the circulation of SLE patients over time and across tissues in a murine model of lupus. The persistence of clonally expanded Tph and Tfh cells may pose a substantial immunologic barrier to the induction of long-lasting immune tolerance with immunosuppressive or B cell-depleting therapies. Targeting these expanded T cell clones directly may be required to fully erase the autoimmune T cell response established early on in patients with SLE.

## Acknowledgements

The work has been supported in part by funding from Merck & Co., Inc (to K.H.C. and D.A.R.), and by a Burroughs Wellcome Fund Career Award in Medical Sciences and National Institute of Arthritis and Musculoskeletal and Skin Diseases, NIH P01 AR070253 (to D.A.R.). P.T.S. was supported by NIH grant R01AI200132. The authors would like to thank Junning Case at the Center for Cellular Profiling at Brigham and Women’s Hospital for cell sorting, and the rest of the CCP team for scRNA-seq data generation. We thank the Accelerating Medicines Partnership: RA/SLE Network for generation of the rheumatoid arthritis synovial scRNA-seq data (syn52297840) and SLE mass cytometry data (syn65922997). Data from the AMP RA/SLE Network were obtained from the ARK Portal (arkportal.synapse.org). We thank the ImmuNexUT (Immune Cell Gene Expression Atlas from the University of Tokyo) for sharing the bulk RNA-seq data for autoimmune diseases.

## Author contributions

T.S. and J.S. conceived the project, performed experiments, analyzed data, and wrote the manuscript. Y.X. analyzed transcriptomic data. R.W., S.S., T.A., K.E.M., Y.G., R.B.R., and P.Y.L. performed experiments. P.T.S. designed the fate mapping model and supervised analyses of this system. A.H. assisted with mass cytometry analysis. Y.Q., M.A.S., S.E.A., and M.C.L. participated in study design. K.F. generated transcriptomic data. K.H.C. and D.A.R. conceived the project, supervised the work, recruited the patients, and wrote the manuscript. All authors discussed the results and commented on the manuscript.

## Methods

### Patient samples

This study was approved by the research ethics committees (BWH IRB #: 2014P002558), and informed consent was obtained from all participating patients for the collection of specimens. All SLE patients met the 1997 ACR classification criteria, the 2012 SLICC criteria, and the 2019 EULAR/ACR classification criteria (*54–56*). For the longitudinal SLE cohort used for TCR and BCR sequencing, 9 SLE patients were included who were within 6 months of disease diagnosis and had not received major immunosuppressive therapies (treatment with prednisone ≤10 mg and hydroxychloroquine was permitted), along with 10 non-inflammatory controls. Samples from the 9 SLE patients were collected at the time of enrollment, 6 months after enrollment, and 1 year after enrollment. For the high-risk for lupus study, patients who did not initially meet any classification criteria for SLE but scored 4-9 points on the EULAR/ACR 2019 classification criteria for SLE were followed longitudinally, and 3 patients who subsequently fulfilled one of the SLE classification criteria during follow-up were studied. PBMC samples were isolated by density gradient centrifugation using Ficoll-Paque Plus (GE Healthcare) and cryopreserved in FBS with 10% DMSO via slow freezing, followed by storage in liquid nitrogen.

### Bulk TCR and BCR sequencing

Frozen PBMC samples were thawed into RPMI Medium 1640 (Life Technologies #11875-085), supplemented with 10% heat-inactivated FBS (Life Technologies #16000044). The samples were stained with flow cytometry antibodies for 30 minutes on ice. After staining, the cells were washed once in PBS with 1% BSA, passed through a 70 μm filter, and stained with 1 ul of a 1:100 diluted Propidium Iodide Solution (BioLegend, #421301). 7 T cell subsets and 4 B cell subsets were directly sorted from the samples using a 5L BD FACSAria Fusion cell sorter into RLT lysis buffer kept on ice. The lysate was then frozen at −80°C until shipment to iRepertoire. Bulk TCRB sequencing data were generated from DNA, and bulk BCR-IGHV sequencing data from RNA. Each lysate was split into two samples to create duplicate data. Read count data for each sample were generated by iRepertoire, including CDR3 sequences, sequence read counts for each clone, and information on the identified V, D, and J genes. These data were subsequently loaded into R (v4.2.0) for further analysis. Non-productive sequences identified by IMGT/V-QUEST were removed, and the count data were analyzed using the Immunarch package. For the TCR overlap analysis, samples were first downsampled to 5,000 clones, and the overlap coefficient between each pair of repertoires was calculated using the repOverlap function in the Immunarch with the "overlap" method.

### Single-Cell RNA and TCR sequencing for human samples

Frozen PBMC samples from 3 out of 9 SLE patients were thawed in RPMI Medium 1640 with 10% FBS. Cells were incubated with an Fc receptor blocking solution, Human TruStain FcX™ (BioLegend), in Cell Staining Buffer (BioLegend) for 10 minutes at 4°C, followed by staining with flow antibodies for 30 minutes on ice. 5 T cell subsets, from non-Treg memory cells, were sorted into RPMI Medium 1640 with 10% FBS. After washing with Cell Staining Buffer, the cells were stained with a hashtag antibody (BioLegend) for 30 minutes on ice. Cells were then washed three times with Cell Staining Buffer, and 30,000 cells from each subset/donor were mixed. Then the cells were stained with TotalSeq™-C Human Universal Cocktail (BioLegend) for 30 minutes on ice, washed three times, and resuspended in 0.4% BSA in PBS. After counting live cells in 0.4% BSA in PBS using Trypan blue, a total of 160,000 cells in two batches were loaded onto a Chromium Next GEM Chip G (10x Genomics). cDNA and library preparation were conducted according to the manufacturer’s protocol. mRNA libraries were sequenced using the NovaSeq S4 flow cell (Illumina), and CITE-seq antibody-derived tag (ADT) libraries were sequenced using the HiSeq X Ten System (Illumina). Unique molecular identifier (UMI) counts for mRNA and ADT were quantified using Cell Ranger v7.1.1. FASTQ files were aligned to the GRCh38 human reference genome using Cell Ranger v7.1.1 (10x Genomics), and gene and ADT reads were quantified simultaneously using the Cell Ranger count.

The files were then processed with the Seurat R package (v5.1.0) (*57*). Expression of each hashing antibody was normalized using the centered-log ratio (CLR) method, followed by demultiplexing with the HTODemux function and a positive quantile of 0.99. Doublets and cells without hashtag staining were excluded. For quality control, we removed cells expressing fewer than 200 genes, more than 3,000 genes, or containing more than 10% of total UMIs associated with mitochondrial genes. After QC, mRNA expression was log-normalized (scale.factor = 10,000), and protein expression was normalized with CLR. The top 2,000 most highly variable genes, selected through variance stabilizing transformation, were scaled. Next, 20 principal components (PCs) were calculated based on mRNA data, and samples were batch-corrected using the Harmony R package. UMAP visualization and clustering (resolution = 0.5) were performed. Tissue Tph/Tfh signatures were calculated using the differentially expressed genes (DEGs) of Tph/Tfh clusters with the AddModuleScore function in Seurat.

Sequencing reads for the TCR library were processed through the Cell Ranger workflow (v7.1.1). Reads were aligned to a TCR reference (vdj-GRCh38), productive contigs were filtered, and CDR3 sequences were identified for each cell. Cells with matching both CDR3alpha and CDR3beta were grouped into the same clonotype. Each cell’s TCR was then paired with its transcriptomic data by matching cell barcodes. These files were then analyzed using the scRepertoire package (*58*). To identify transcription factors potentially driving the C5 transcriptional program, TF activities were inferred using decoupleR (*59*) based on the signed TF–target interactions in the CollecTRI-derived regulons (*60*). For the tissue Tph/Tfh signature, AMP clusters 1, 4, and 7 were compared with all other CD4 T-cell clusters using Seurat FindMarkers with the Wilcoxon rank-sum test, min.pct = 0.25, and logfc.threshold = 0.25. To identify genes specifically enriched in Tph/Tfh cells, differentially expressed genes associated with AMP cluster 3, corresponding to Treg cells, were separately identified using the same approach and excluded from the Tph/Tfh gene set. The remaining genes were ranked by average log₂ fold change, and the top 100 positively enriched genes were selected.

### Mass cytometry

Cryopreserved PBMC were thawed into RPMI Medium 1640 (Life Technologies #11875-085) supplemented with 5% heat-inactivated fetal bovine serum (Life Technologies #16000044), 1 mM GlutaMAX (Life Technologies #35050079), antibiotic-antimycotic (Life Technologies #15240062), 2 mM MEM non-essential amino acids (Life Technologies #11140050), 10 mM HEPES (Life Technologies #15630080), 2.5 x 10^-5^ M 2-mercaptoethanol (Sigma-Aldrich #M3148), 20 units/mL sodium heparin (Sigma-Aldrich #H3393), and 25 units/mL benzonase nuclease (Sigma-Aldrich #E1014). Cells were counted and 0.5-1×10^6^ cells from each sample were transferred to a polypropylene plate for staining. The samples were spun down and aspirated. 5 μM of cisplatin viability staining reagent (Fluidigm #201064) was added for two minutes and then diluted with culture media. After centrifugation, Human TruStain FcX Fc receptor blocking reagent (BioLegend #422302) was used at a 1:100 dilution in PBS with 2.5 g bovine serum albumin (Sigma Aldrich #A3059) and 100 mg of sodium azide (Sigma Aldrich #71289) for 10 minutes followed by incubation with conjugated surface antibodies for 30 minutes. All antibodies were obtained from the Harvard Medical Area CyTOF Antibody Resource and Core (Boston, MA). 16% stock paraformaldehyde (Fisher Scientific #O4042-500) dissolved in PBS was used at a final concentration of 4% formaldehyde for 10 minutes in order to fix the samples before permeabilization with the FoxP3/Transcription Factor Staining Buffer Set (ThermoFisher Scientific #00-5523-00). The samples were incubated with SCN-EDTA coupled palladium based barcoding reagents for 15 minutes and then combined into a single sample. Conjugated intracellular antibodies were added into each tube and incubated for 30 minutes. Cells were then fixed with 1.6% formaldehyde for 10 minutes. DNA was labelled for 20 minutes with an 18.75 μM iridium intercalator solution (Fluidigm #201192B). Samples were subsequently washed and reconstituted in Milli-Q filtered distilled water in the presence of EQ Four Element Calibration beads (Fluidigm #201078) at a final concentration of 1×10^6^ cells/mL. Samples were acquired on a Helios CyTOF Mass Cytometer (Fluidigm). Cytometry data were normalized and debarcoded. Live cells were gated as DNA^+^ Live^+^ Beads^-^ using FlowJo (BD). Batch-corrected total live cells were then analyzed using the Seurat R package (v5.1.0) (*57*) for PCA, neighborhood analysis, and clustering visualized by UMAP. Cells associated with SLE were identified using CNA (*32*).

To independently validate our findings, we additionally reanalyzed previously published CyTOF data from patients with biopsy-proven lupus nephritis (LN) and healthy controls reported by Horisberger et al. (*7*). Processed mass cytometry data and corresponding clinical metadata from the AMP RA/SLE phase II cohort were obtained through the ARK Portal (Synapse accession syn65922997) and analyzed separately from the cohort described above. For analyses according to renal treatment response, patients with a baseline urine protein-to-creatinine ratio (UPCR) ≥1 g/g were classified based on renal response at week 52 according to the criteria used in the original study. Complete response (CR) was defined as UPCR <0.5 g/g, serum creatinine ≤1.3 mg/dL or, if abnormal at baseline, <125% of the baseline value, and prednisone <10 mg/day. No response (NR) was defined as failure to meet the criteria for either complete or partial response. Comparisons between CR and NR were performed using patients meeting these respective response definitions.

### Mice and pristane treatment

C57BL/6J, *Bcl6*^fl/fl^, CD4-Cre, B6.Cg-Sle1^NZM2410/Aeg^ Yaa/DcrJ, and Rosa26-Lox-STOP-Lox-YFP mice were obtained from The Jackson Laboratory. To generate mice with T-cell-specific deletion of Bcl6, *Bcl6*^fl/fl^ mice were crossed with CD4-Cre hemizygous mice. Sex-matched *Bcl6*^fl/fl^ CD4Cre⁺ mice and Cre-negative littermate controls were used for experiments.

The constitutive *Cxcr5-Cre* mouse was generated by inserting a Cre recombinase sequence downstream of exon 2 of the endogenous *Cxcr5* locus through a P2A linker using CRISPR/Cas9 targeting in C57BL/6 embryonic stem cells. *Cxcr5-Cre* mice were crossed with Rosa26-Lox-STOP-Lox-YFP reporter mice to generate *Cxcr5-Cre Rosa26-LSL-YFP* mice. In these animals, current or previous expression of *Cxcr5* results in irreversible Cre-mediated excision of the STOP cassette and permanent YFP labelling.

Pristane-induced lupus was initiated by a single intraperitoneal injection of 0.5 ml pristane (Sigma-Aldrich). Mice were maintained for the indicated experimental periods and monitored weekly. All animal procedures were approved by the Brigham and Women’s Hospital Institutional Animal Care and Use Committee under protocol 2018N000053.

### Mouse tissue processing and flow cytometry

Mice were euthanized at the indicated time points, and tissues were processed immediately. Spleen, lymph nodes, granulomas, and lungs were mechanically dissociated to generate single-cell suspensions. Lung tissue was additionally digested in RPMI containing 100 µg/ml Liberase TL and 100 µg/ml DNase I (both Sigma-Aldrich) for 30 min at 37°C with agitation. Digestion was quenched with excess medium, and cells were washed and passed through a 70-µm filter. Lung cells were subsequently incubated for a further 20 min at 37°C to allow recovery of cell-surface CXCR5 expression.

Cells were incubated with Live/Dead Aqua viability dye (Thermo Fisher Scientific) and Fc receptor-blocking antibody (BioLegend) in PBS for 10 min at 4°C. Cells were then stained using standard procedures with TotalSeq hashing antibodies, CITE-seq antibodies against ICOS, I-A/I-E, CD200, CXCR6, CCR2, and CX3CR1, and a flow-cytometry panel containing antibodies against CD45.2, B220, CD4, TCRβ, CD44, CD62L, CD25, IL-7Rα, PD-1, and CXCR5 (BioLegend).

For experiments using *Cxcr5-Cre Rosa26-LSL-YFP* mice, splenocytes from four mice were pooled before sorting. Cells were gated sequentially as singlets, live CD45.2⁺TCRβ⁺CD4⁺ cells. Four mutually exclusive populations were isolated according to YFP, CXCR5, and PD-1 expression: CXCR5⁺YFP⁺, CXCR5⁻YFP⁺, YFP⁻CXCR5⁻PD-1⁺, and YFP⁻CXCR5⁻PD-1⁻ cells. The latter two populations are referred to as PD-1⁺ and PD-1⁻, respectively. The original sorting identity was retained in the single-cell metadata, allowing transcriptionally defined populations to be related to their CXCR5-expression and lineage-labelling histories.

### Bulk RNA sequencing

For bulk RNA sequencing, 2,000 cells from each population were sorted directly into TCL lysis buffer (Qiagen) containing 1% β-mercaptoethanol (Sigma-Aldrich). Lysates were vortexed to ensure complete lysis and snap-frozen on dry ice. Libraries were generated and sequenced using the SMART-Seq2 protocol at the Broad Institute. RNA-seq reads were quantified against the Mus musculus Ensembl release 104 transcriptome based on the GRCm39 genome assembly using Salmon v1.4.0.

Salmon quantification files were imported into R using tximeta, and sample metadata were generated according to mouse, tissue, and sorted cell population. Raw counts were normalized and analysed using DESeq2. Lowly expressed genes and mitochondrial, olfactory-receptor, and unannotated genes were excluded before downstream analysis. Where indicated, mouse-associated batch effects were removed using limma. Principal-component analysis was performed using prcomp, and results were visualized using ggplot2. Differential expression was assessed using DESeq2. Variance-stabilizing transformed expression values were used for heatmaps and hierarchical clustering with pheatmap.

### Mouse single-cell RNA sequencing and preprocessing

For single-cell sequencing, total CD4 T cells or the indicated flow-sorted populations were collected into PBS containing 0.4% BSA. For the 12-month and *Bcl6* conditional-knockout experiments, equal numbers of cells from each mouse or genotype were counted, pooled, and processed using the 10x Genomics Chromium platform at the Brigham and Women’s Hospital Center for Cellular Profiling. Approximately 30,000 cells were loaded for the 12-month experiment (*n* = 5 mice), and approximately 100,000 cells were loaded for the *Bcl6*^fl/fl^ CD4Cre⁻/⁺ experiment (*n* = 5 mice per group).

For the CXCR5-lineage-reporter experiment, splenocytes from four *Cxcr5-Cre Rosa26-LSL-YFP* mice were pooled before sorting. A total of 5,000 cells was collected from each of the four CXCR5⁺YFP⁺, CXCR5⁻YFP⁺, PD-1⁺, and PD-1⁻ populations. Each sorted population was labelled with a distinct oligonucleotide-conjugated multiplexing antibody (on chip multiplexing), and the four populations were pooled to produce an input of approximately 20,000 cells for 10x Genomics Chromium sequencing.

Raw sequencing data from the 12-month and 3-month pristane datasets were processed using Cell Ranger v6.1.1 and aligned to the 10x Genomics mm10-2020-A mouse reference transcriptome. Data from the CXCR5-reporter mouse experiment were processed using Cell Ranger v9.0.1 and aligned to the 10x Genomics mm10-3.0.0 reference transcriptome. Both references are based on the GRCm38/mm10 mouse genome assembly. Gene-expression matrices were imported into R and analysed using Seurat v5.5.1.

Hashtag oligonucleotide or oligonucleotide-conjugated multiplexing counts were normalized, and cells were assigned to their sample or sorted population of origin using HTODemux. Cells classified as hashtag-negative or doublets were excluded. Cells expressing fewer than 300 genes or containing more than 12% mitochondrial transcripts were also excluded.

After quality control, the 12-month dataset contained 20,794 splenic and 16,793 lung CD4 T cells. The three-month control dataset contained 33,554 splenic and 24,923 lung CD4 T cells, while the three-month *Bcl6*^fl/fl^ CD4Cre⁺ dataset contained 33,791 splenic and 23,051 lung CD4 T cells. The CXCR5-lineage-reporter dataset contained 11,308 splenic CD4 T cells, comprising 2,662 CXCR5⁻YFP⁺ cells, 2,648 CXCR5⁺YFP⁺ cells, 3,104 PD-1⁻ cells, and 2,894 PD-1⁺ cells.

Counts were normalized using NormalizeData and scaled using ScaleData. The 2,000 most variable genes were identified using FindVariableFeatures, followed by principal-component analysis. Harmony was used to account for mouse- or cohort-associated batch effects where applicable. UMAP dimensionality reduction and graph-based clustering were performed using the indicated number of harmonized principal components and clustering resolution for each analysis. DimPlot, FeaturePlot, and DotPlot were used for visualization. Differential expression between clusters or subclusters was assessed using FindMarkers with the Wilcoxon rank-sum test.

### Construction of bulk RNA-sequencing Tph and Tfh signatures

Tissue-specific Tph and Tfh signatures were generated from the bulk RNA-sequencing data using naïve CD4 T cells as the reference population. DESeq2 contrasts were performed separately for splenic Tph versus splenic naïve, splenic Tfh versus splenic naïve, lung Tph versus lung naïve, and lung Tfh versus lung naïve cells. Independent filtering was disabled when extracting DESeq2 results.

For each comparison, genes with an adjusted *P* value below 0.05 and a positive log₂ fold change were ranked in descending order of log₂ fold change, and the top 100 genes were selected. These naïve-referenced lists were then filtered using the corresponding direct Tph-versus-Tfh comparison. Genes retained in the Tph signature were upregulated in Tph relative to naïve cells and were either higher in Tph than Tfh cells or showed no evidence of preferential expression in Tfh cells. The reciprocal criteria were applied to the Tfh signature. Genes with expression patterns inconsistent with the intended Tph or Tfh identity were removed, and overlapping genes were excluded to generate non-overlapping signatures.

Signatures were generated separately for the spleen and lung to preserve tissue-associated transcriptional programmes. This strategy used the naïve population to identify genes acquired during differentiation into each B-cell-helper state, while the direct Tph-versus-Tfh comparison ensured that the resulting signatures retained the appropriate direction of expression between the two related populations.

### Mouse B-cell-helper, Tph, and Tfh signatures

A broad B-cell-helper (BCH) signature was generated from the 12-month scRNA-seq dataset. The integrated dataset was divided by tissue, and differential-expression analysis was performed separately within the spleen and lung using Seurat FindMarkers. Splenic BCH cluster 2 was compared with all other splenic CD4 T cells, while lung BCH cluster 4 was compared with all other lung CD4 T cells. Only positively enriched genes with an adjusted *P* value below 0.05 were retained. Genes were ranked by average log₂ fold change, and the top 50 genes from each comparison were used to generate tissue-specific spleen and lung BCH signatures.

Tph- and Tfh-specific signatures were generated by directly comparing the transcriptionally defined splenic Tph and Tfh subclusters from the same 12-month dataset using FindMarkers. The 50 genes with the highest positive average log₂ fold change in Tph versus Tfh cells were used as the mouse Tph signature. The reciprocal comparison was used to select the top 50 genes for the mouse Tfh signature. These signatures were subsequently used to identify corresponding BCH, Tph, and Tfh populations in the independent three-month *Bcl6* conditional-knockout dataset.

### Human-derived Tph and Tfh signatures

Human-derived Tph and Tfh signatures were generated from published single-cell RNA-sequencing datasets of synovial CD4 T cells from the Accelerating Medicines Partnership RA cohort and human tonsillar CD4 T cells, respectively. For the human Tph signature, the Tph clusters were compared with all other CD4 T-cell clusters using Seurat FindMarkers with the Wilcoxon rank-sum test, min.pct = 0.25, and logfc.threshold = 0.25. Genes were ranked by average log₂ fold change, and the top 50 positively enriched genes were selected. For the human Tfh signature, Tfh cluster was compared with all other tonsillar CD4 T-cell clusters using the same parameters, and the top 50 positively enriched genes were selected. Human genes were converted to their mouse orthologues using homologene, with human taxonomy identifier 9606 and mouse taxonomy identifier 10090. Genes without an identifiable mouse orthologue were excluded, and duplicated mouse gene symbols were collapsed. The resulting gene sets were used to score the mouse single-cell and pseudobulk datasets.

An additional Tph signature was derived from the human peripheral-blood Tph population (C5) identified in this study. C5 was compared with all other clusters using Seurat FindMarkers with the Wilcoxon rank-sum test, only.pos = TRUE, min.pct = 0.25, and logfc.threshold = 0.75. Up to 400 positively enriched genes were selected in descending order of average log₂ fold change. Genes shared with an analogous signature derived from the human Tfh population (C0) were removed. Human genes were converted to mouse orthologues using homologene, and any overlap introduced by orthologue mapping was also removed. The resulting C5-derived Tph signature was used for mouse-level GSVA analysis.

### GSVA and gene-signature analysis

Single-cell signature scores were calculated using UCell (version 2.16), added to the corresponding Seurat metadata, and visualized using feature and violin plots. For mouse-level GSVA, raw counts were aggregated for each mouse–subcluster combination using Seurat AggregateExpression, normalized using the trimmed mean of M-values method in edgeR, and converted to log₂ counts per million. GSVA was performed separately for spleen and lung using the C5-derived Tph signature and a Gaussian kernel.

### T-cell receptor sequencing and clonotype analysis

TCR libraries were processed using Cell Ranger V(D)J versions 6.1.1 or 9.0.1, according to the dataset. Assembled TCR sequences were integrated with the corresponding Seurat objects using scRepertoire version 1.7.2. Cells lacking paired TCRα and TCRβ chains were excluded. Clonotypes were defined by identical TCR V(D)J gene usage and identical CDR3 nucleotide sequences across the paired α and β chains.

Clonal-expansion categories were defined using the default scRepertoire thresholds: single, one cell; small, two to five cells; medium, six to 20 cells; large, 21 to 100 cells; and hyperexpanded, 101 to 500 cells. Expanded clonotypes were defined as clonotypes detected in at least two cells. Clonotype expansion and distribution were visualized on UMAP embeddings using scRepertoire, Seurat, and ggplot2.

Clonotype sharing among clusters, subclusters, tissues, and sorted populations was quantified by extracting expanded clonotypes from each population and identifying matching paired TCR sequences in comparison populations. UpSet plots were used to visualize clonotype intersections. Morisita indices were calculated using scRepertoire to quantify repertoire similarity while accounting for clone abundance.

For the CXCR5-lineage-reporter experiment, each cell’s original sorting identity was retained after integration with the scRNA-seq data. Clonotypes were classified as PD-1⁺ only when cells belonging to the clonotype were detected exclusively in the PD-1⁺ sorting compartment, YFP⁺ only when they were detected exclusively in one or both YFP⁺ compartments, or PD-1⁺ and YFP⁺ when members of the same clonotype were recovered from both compartments. This analysis allowed clonally related transcriptional states to be connected to their CXCR5-lineage histories. As cells from the four reporter mice were pooled before sorting, clonotypes in this experiment could not be assigned to individual animals. However, analysis of independently tracked mice in the separate 3- and 12-month pristane datasets showed that inter-mouse clonotype sharing was exceptionally rare (data not shown). Thus, although public clonotypes cannot be completely excluded from the pooled reporter experiment, they are unlikely to account for a substantial proportion of the observed sharing between cell populations.

### SCENIC regulon analysis

Transcription-factor regulons were inferred using SCENIC version 1.3.1 from all cells in clusters 1 and 3 of the CXCR5-lineage-reporter dataset. Analyses were performed using AUCell version 1.32.0, RcisTarget version 1.29.0, and GENIE3 version 1.32.0.

Gene regulatory networks were initially inferred using GENIE3. Transcription factor–target relationships were refined by motif-enrichment analysis using RcisTarget and the mouse mm9 500bp-upstream-7species.mc9nr and tss-centered-10kb-7species.mc9nr databases. For visualization of the Bcl6 regulon in Figure 7K, the AUC score for each cell was averaged with those of its 30 nearest neighbours in the two-dimensional UMAP embedding, identified using FNN::get.knn. These smoothed values were used to colour cells in the plot; the underlying AUCell scores were not altered.

### Covarying neighborhood analysis

Differences in cellular representation between *Bcl6*^fl/fl^ CD4Cre⁻ and *Bcl6*^fl/fl^ CD4Cre⁺ mice were evaluated using covarying neighborhood analysis (rcna v0.0.99). The analysis was performed on the integrated CD4 T-cell object using the previously constructed RNA nearest-neighbor graph (RNA_nn), with genotype as the tested biological variable and mouse as the experimental unit. Genotype was encoded as *Bcl6*^fl/fl^ CD4Cre⁻= 1 and *Bcl6*^fl/fl^ CD4Cre⁺ = 0. No additional batch variables or covariates were included. Normalized neighborhood correlation scores (cna_ncorrs) were projected onto the UMAP embedding, with positive and negative values indicating neighborhoods preferentially represented by control and Bcl6-deficient cells, respectively.

### In vitro assessment of CXCR5 downregulation

Splenic Tfh cells were isolated from B6.Sle1^yaa^ mice by magnetic enrichment of CD4 T cells (Miltenyi Biotec), followed by sorting of live B220⁻CD4⁺GITR⁻CXCR5⁺PD-1⁺ cells. Sorted Tfh cells were cocultured with autologous B cells at a 1:1 ratio in the presence of plate-bound anti-CD3 and soluble anti-IgM to provide TCR and BCR stimulation, respectively.

Cultures were maintained for five days in complete RPMI 1640 containing 10% FBS, 100 U/ml penicillin, 100 µg/ml streptomycin, 2 mM L-glutamine, and 50 µM β-mercaptoethanol. Cells were harvested at the end of the culture period and analysed by flow cytometry. Live CD4⁺I-A⁺ cells were assessed for PD-1 and CXCR5 expression to quantify the emergence of a CXCR5⁻PD-1^hi^ phenotype.

### ELISA

Serum total IgG and anti-dsDNA IgG concentrations were measured in *Bcl6*^fl/fl^ CD4Cre⁻ and *Bcl6*^fl/fl^ CD4Cre⁺ mice three months after pristane treatment using commercially available ELISA kits (Chondrex) according to the manufacturer’s instructions. Differences between genotypes were assessed using a two-sided Mann–Whitney U test.

---

**Extended Data Figure 1.**
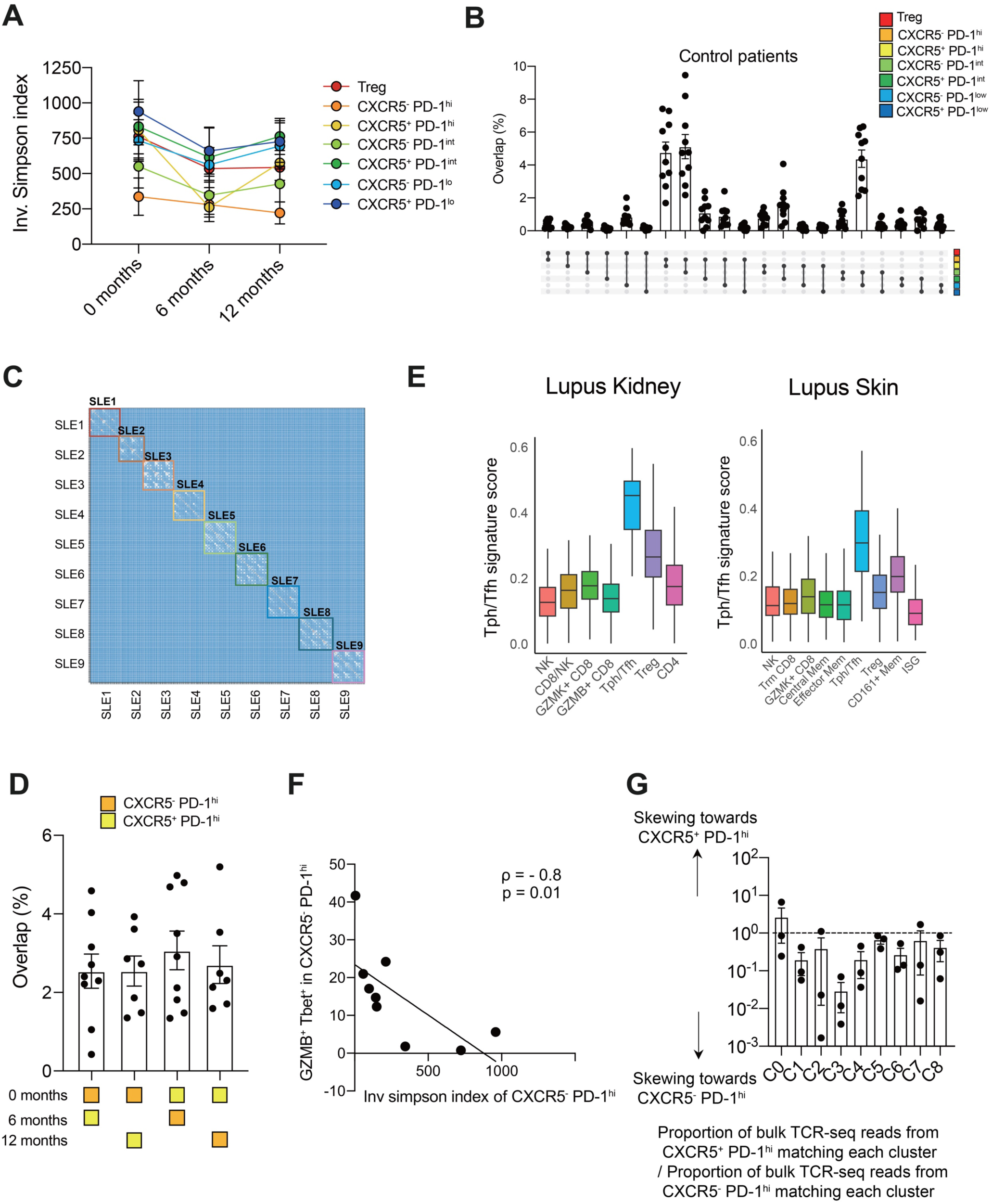
TCR repertoire features in SLE patients. **A.** TCR clonality changes over time (SLE: n = 9). TCR clonality was assessed by inverse Simpson Index (downsampled to 5000 reads). **B.** TCR repertoire overlap between Tph cells, Tfh cells, Tregs, and 4 other comparator CD4 T cell subsets in non-inflammatory controls (n = 10, downsampled to 5000 reads). **C.** Heatmap of TCR repertoire overlap based on CDR3 beta similarity across all samples from longitudinal SLE patients (n = 378 samples). **D.** Frequency of unique TCR repertoire to Tph or Tfh cells at baseline detected at 6 months or 1 year (Time B : n = 9, Time C : n = 7, downsampled to 5000 reads). **E.** Tissue Tph/Tfh score of lupus kidney scRNAseq clusters (*30*) and lupus skin scRNAseq clusters (*31*). **F.** Correlation analysis between Inverse Simpson Index (shown in Figure 1B) and frequencies of GZMB^+^ Tbet^+^ cells in CXCR5^-^ PD-1^hi^ Tph cells using 9 baseline SLE samples. **G.** The ratio of the proportion of CXCR5^+^ PD-1^hi^ bulk TCR-seq reads matching scTCRseq in each cluster to the proportion of CXCR5^-^ PD-1^hi^ bulk TCR-seq reads matching scTCRseq in each cluster (n = 3).

**Extended Data Figure 2.**
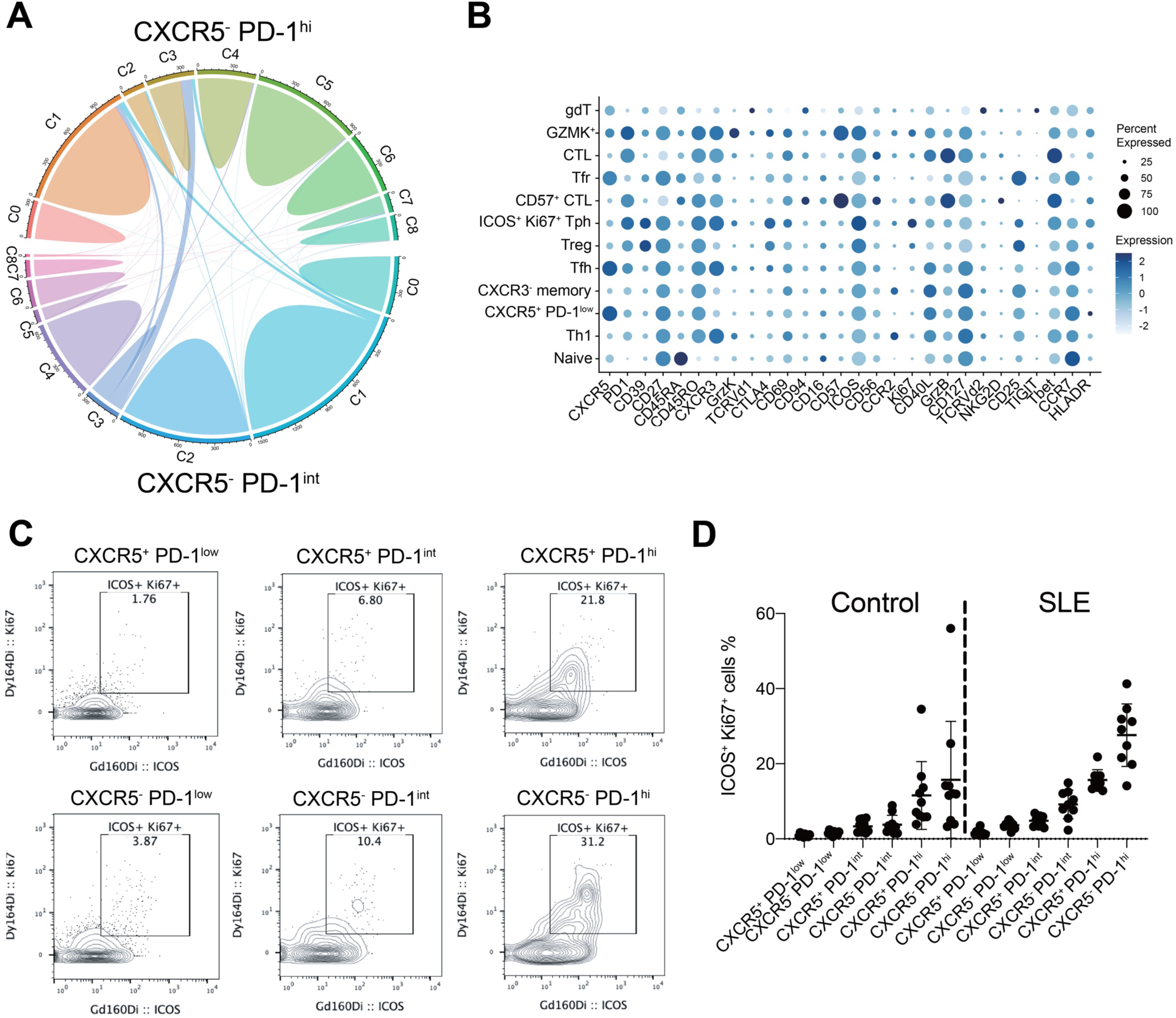
TCR overlap and mass cytometric validation of C5 detected by scRNAseq. **A**. Circos plot showing the TCR sequence overlap in each cluster between CXCR5^-^ PD-1^hi^ cells and CXCR5^-^ PD-1^int^ cells. **B.** Marker expression of each cluster in the UMAP. **C**. ICOS^+^ Ki67^+^ cells in CXCR5^-^ PD-1^hi^ Tph, CXCR5^+^ PD-1^hi^ Tfh, CXCR5^-^ PD-1^int^, CXCR5^+^ PD-1^int^, CXCR5^-^ PD-1^low^, CXCR5^+^ PD-1^low^ cells. **D.** Frequencies of **C** in 10 non-inflammatory controls and baseline 9 SLE samples.

**Extended Data Figure 3.**
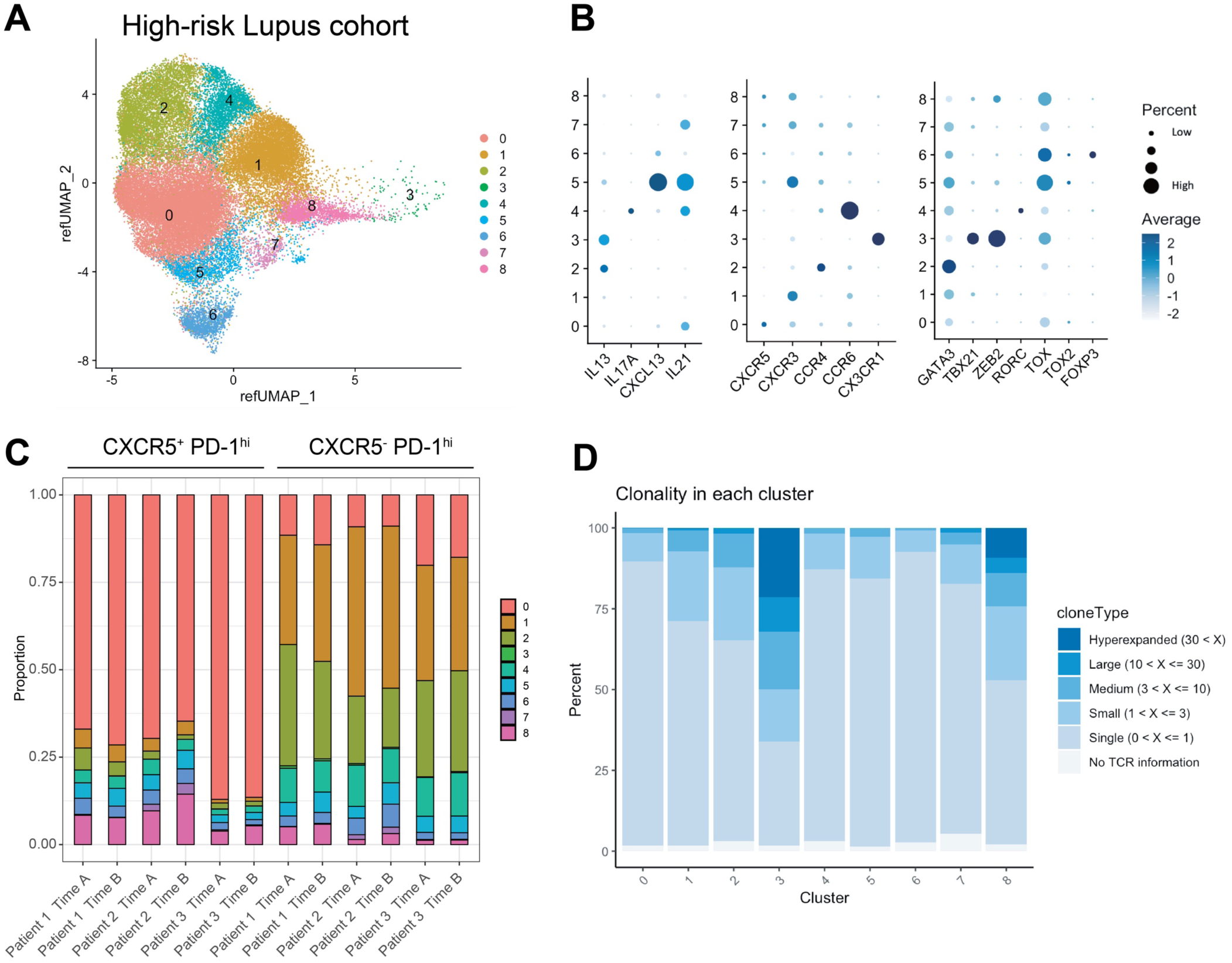
scRNAseq analysis of the high-risk Lupus cohort. **A.** UMAP of CXCR5^-^ PD-1^hi^ cells and CXCR5^+^ PD-1^hi^ cells from the high-risk for lupus cohort. **B.** Gene expression profiles of each cluster in the high-risk for lupus scRNA-seq data. **C.** Cluster abundances in CXCR5^-^ PD-1^hi^ cells and CXCR5^+^ PD-1^hi^ cells from each sample. **D.** Clonality of each cluster in high-risk Lupus PBMC scRNAseq data.

**Extended Data Figure 4.**
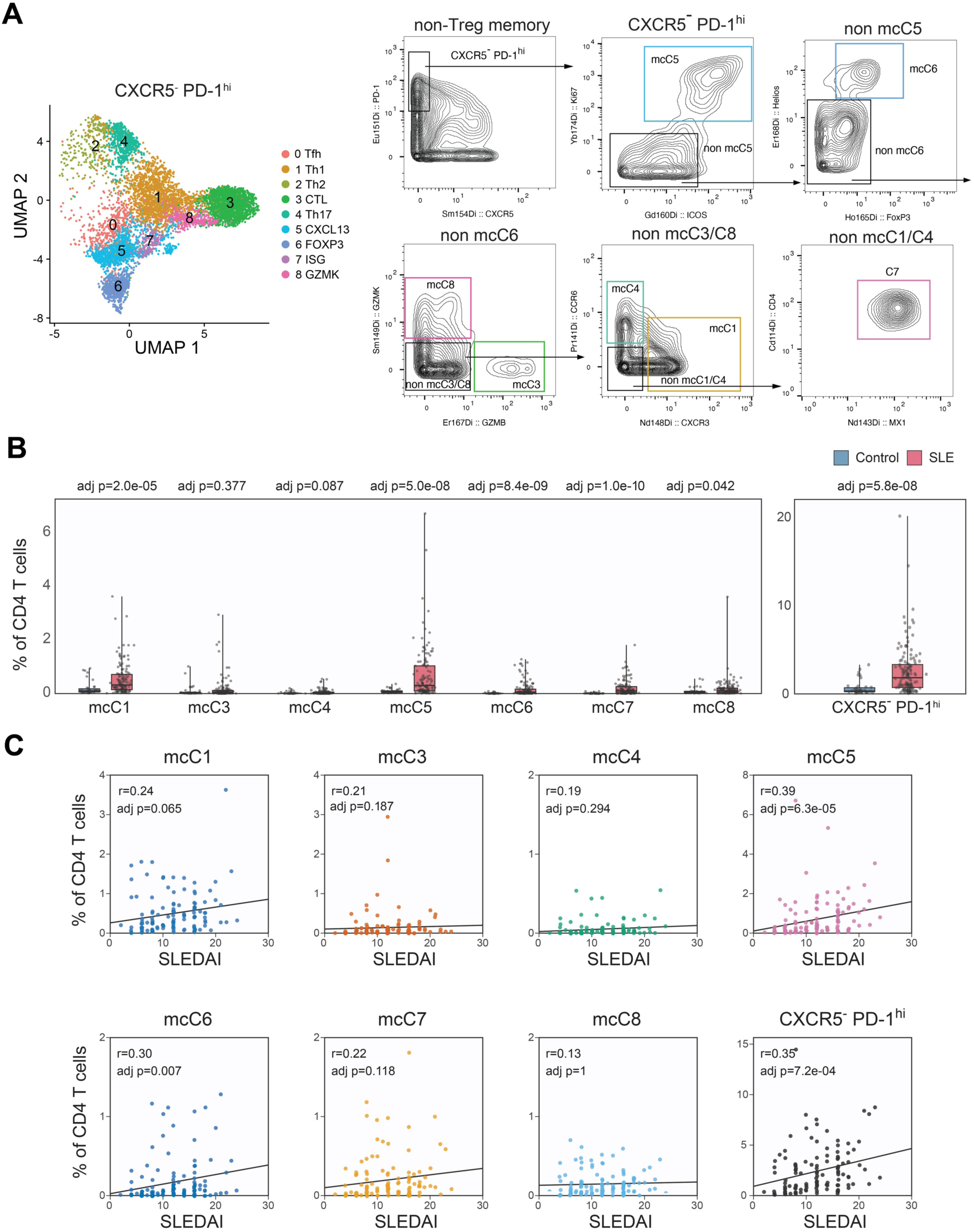
Mass cytometric validation of Tph subpopulations and associations with SLE disease activity. **A.** Representative CyTOF gating strategy for identification of subclusters in CXCR5^-^ PD-1^hi^ cells. **B.** Frequencies of CXCR5^-^ PD-1^hi^ cells and subclusters in healthy controls (HC, n = 40) and patients with SLE (n = 134) from the AMP Phase 2 SLE cohort. P values were adjusted using the Bonferroni correction. **C.** Correlation between the frequencies of CXCR5^-^ PD-1^hi^ cells and subclusters and SLE disease activity (SLEDAI) in the AMP Phase 2 SLE cohort (SLE, n = 134). Spearman’s correlation coefficients are shown, and P values were adjusted using the Bonferroni correction.

**Extended Data Figure 5.**
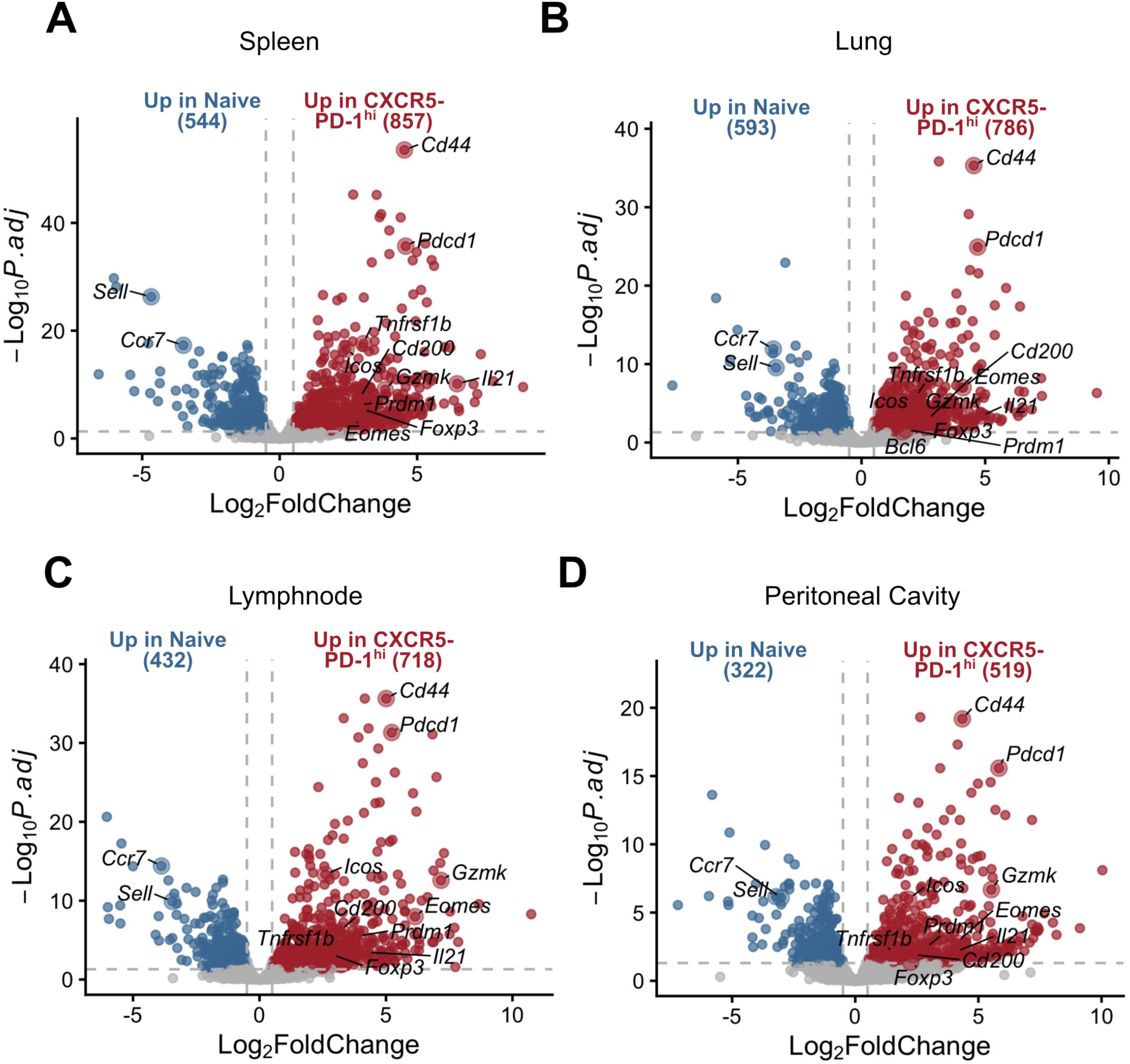
Differential gene expression analysis between CXCR5- PD-1^hi^ and naïve T cells across different tissues. **A.** Volcano plot of differentially expressed genes between spleen naïve CD4 T cells and splenic CXCR5- PD-1^hi^ CD4 T cells. Several significant DEGs are annotated relating to naïve, Th1, Treg and B cell helper function. **B.** Volcano plot of differentially expressed genes between lung naïve CD4 T cells and lung CXCR5- PD-1^hi^ CD4 T cells. Several significant DEGs are annotated relating to naïve, Th1, Treg and B cell helper function. **C.** Volcano plot of differentially expressed genes between lymph node naïve CD4 T cells and lymph node CXCR5- PD-1^hi^ CD4 T cells. Several significant DEGs are annotated relating to naïve, Th1, Treg and B cell helper function. **D.** Volcano plot of differentially expressed genes between granuloma naïve CD4 T cells and granuloma CXCR5- PD-1^hi^ CD4 T cells. Several significant DEGs are annotated relating to naïve, Th1, Treg and B cell helper function.

**Extended Data Figure 6.**
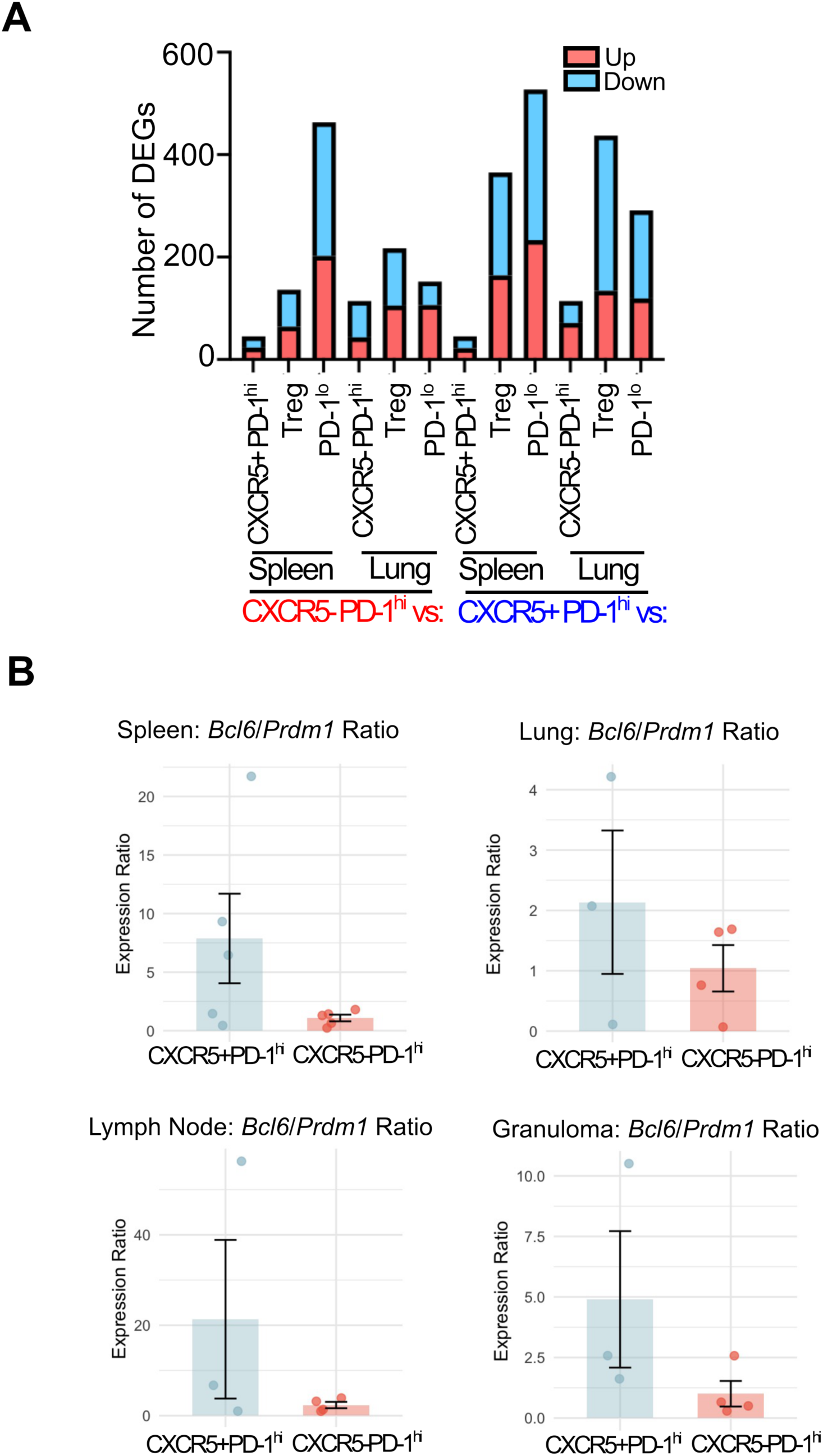
Transcriptional profiles associated with Tfh and Tph cells across different tissues. **A.** Number of differentially expressed genes (DEGs) for each pairwise comparison across spleen and lung. Genes upregulated in CXCR5⁺ PD-1^hi^ (Tfh) cells are shown in red, and genes upregulated in CXCR5⁻ PD- 1^hi^ (Tph) cells are shown in blue. **B.** Ratio of Bcl6 / Prdm1 in Tfh and Tph cells in spleen, lung, lymph node, and granuloma. Normalized counts are shown as mean ± SD for 3–5 mice per group.

**Extended Data Figure 7.**
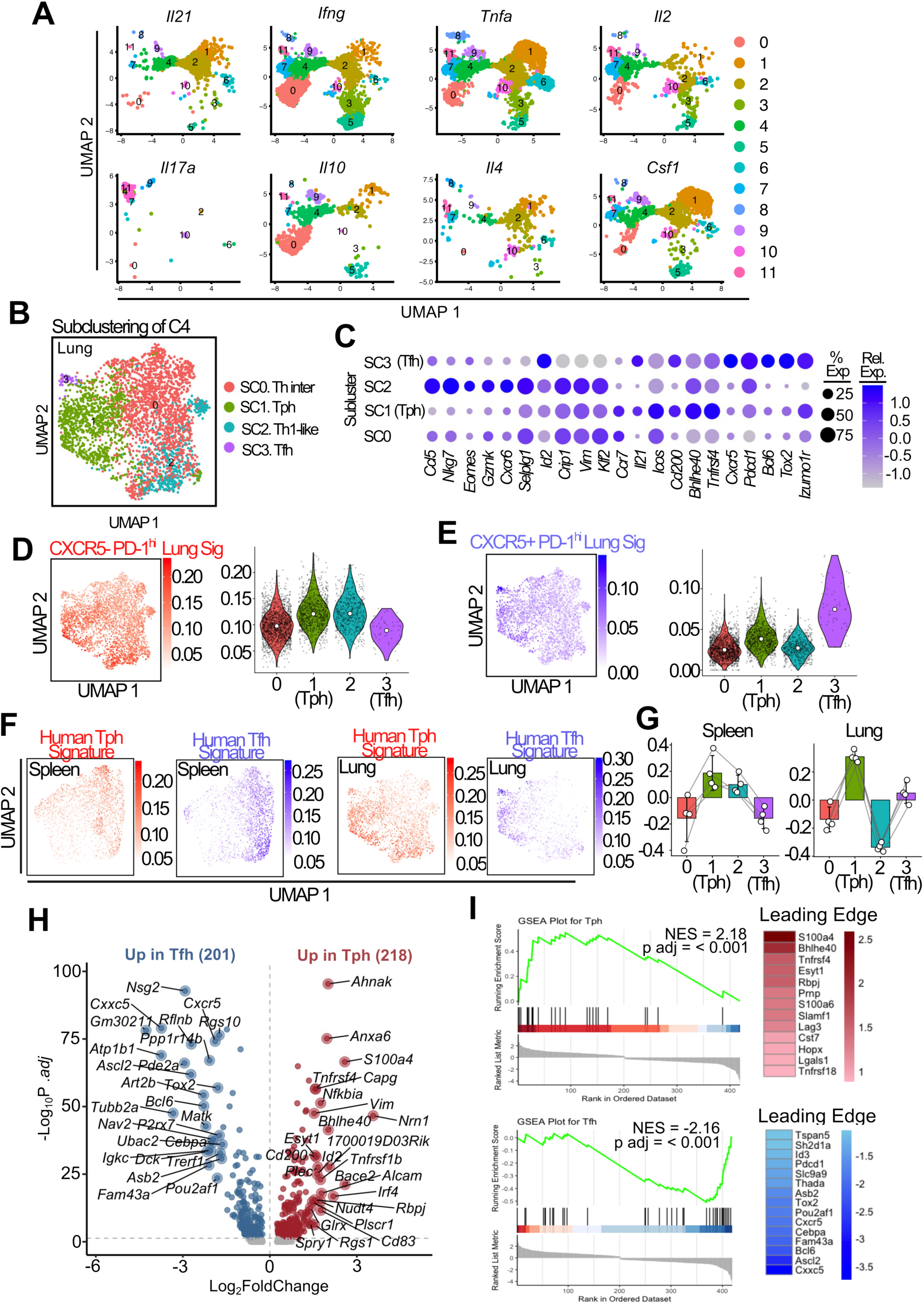
Transcriptional and gene-signature analyses distinguish Tph and Tfh states in murine spleen and lung. **A.** UMAP projections showing the cluster distribution of CD4 T cells expressing the indicated cytokine genes (*Il21, Ifng, Tnfa, Il2, Il17a, Il10, Il4* and *Csf1*). Cells are coloured according to their original CD4 T-cell cluster. **B.** UMAP visualization of subclusters identified following reclustering of lung B-cell-helper cluster 4, annotated as T-helper intermediate (SC0), Tph (SC1), Th1-like (SC2) and Tfh (SC3) populations. **C.** Dot plot showing expression of selected T-cell-state and B-cell-helper genes across the four lung subclusters. Dot size represents the percentage of cells expressing each gene, and colour indicates scaled average expression. **D, E.** UMAP and violin plots showing module scores across the lung subclusters using gene signatures derived from bulk-sorted CXCR5⁻PD-1^hi^ (**D**) or CXCR5⁺PD-1^hi^ (**E**) lung populations. White points indicate the median score within each subcluster. **F.** UMAP projections showing module scores generated using human-derived Tph and Tfh signatures across the splenic and lung B-cell-helper subclusters. **G.** Per-mouse GSVA scores for the human-derived C5 Tph signature across splenic and lung subclusters. Points represent individual mice, with observations from the same mouse connected by lines. **H.** Differential gene-expression analysis comparing splenic Tph (SC1) and Tfh (SC3) cells. Genes significantly upregulated in Tfh and Tph cells are shown in blue and red, respectively, with selected genes annotated. The numbers of differentially expressed genes in each direction are shown above the plot. **I.** GSEA of the lung Tph-versus-Tfh differential-expression ranking using human-derived Tph and Tfh gene signatures. Normalized enrichment scores (NES) and adjusted *P* values are shown. Heatmaps on the right display genes contributing to the leading edge of each enrichment.

**Extended Data Figure 8.**
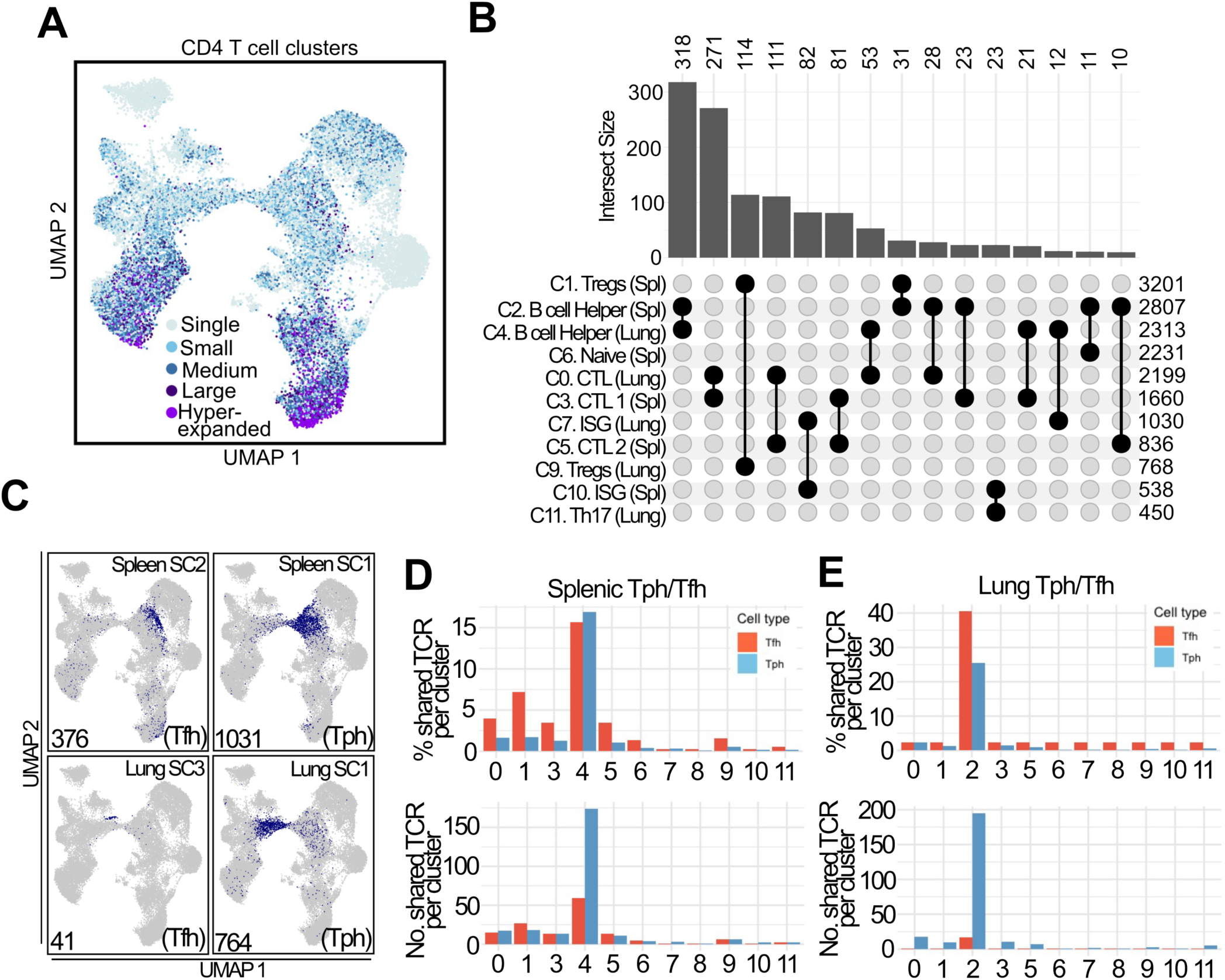
TCR repertoire analysis reveals clonal expansion and clonotype sharing among Tph, Tfh and other CD4 T-cell populations. **A.** UMAP visualization of integrated splenic and lung CD4 T cells coloured according to clonotype expansion category, ranging from single to hyperexpanded clonotypes. **B.** UpSet plot showing TCR clonotype intersections among the indicated CD4 T-cell clusters. Values above the vertical bars indicate the number of clonotypes in each intersection, while values on the right indicate the total number of clonotypes detected in each cluster. **C.** UMAP projections showing the locations of splenic Tfh (SC2) and Tph (SC1) clones and lung Tfh (SC3) and Tph (SC1) clones within the integrated CD4 T-cell dataset. Values indicate the number of cells in each population. **D.** Distribution of clonotypes from splenic Tph and Tfh populations across the original CD4 T-cell clusters. The upper panel shows the percentage of shared clonotypes associated with each cluster, and the lower panel shows the corresponding number of shared clonotypes. **E.** Distribution of clonotypes from lung Tph and Tfh populations across the original CD4 T-cell clusters, presented as the percentage (upper) and number (lower) of shared clonotypes associated with each cluster. Tph and Tfh populations are shown in blue and red, respectively.

**Extended Data Figure 9.**
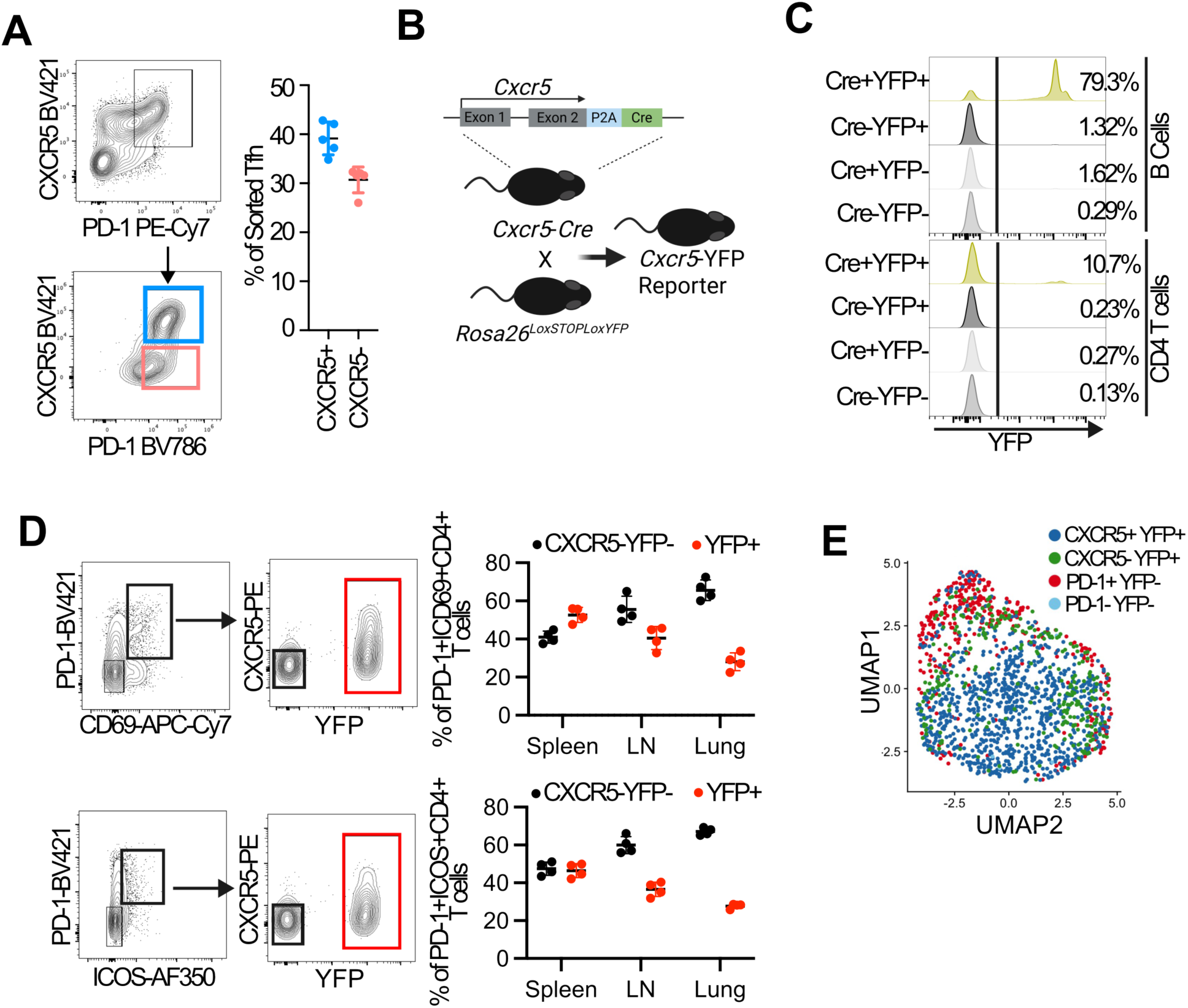
Validation of the CXCR5 lineage reporter and characterization of sorted splenic CD4 T cells. **A.** Representative flow-cytometric gating and quantification of CXCR5 expression after sorted CXCR5⁺PD- 1^hi^ Tfh cells were stimulated in vitro for 5 days. Points represent independent biological samples (*n* = 5); bars show the mean with error bars. **B.** Schematic of the *Cxcr5-Cre Rosa26-LSL-YFP* lineage-reporter model, in which *Cxcr5* expression induces permanent YFP labelling. **C.** YFP expression in B cells and CD4 T cells from reporter and genetic-control mice, showing labelling in mice carrying both the Cre and reporter alleles. **D.** Representative gating of PD-1⁺CD69⁺ and PD-1⁺ICOS⁺ CD4 T cells, with quantification of CXCR5⁻YFP⁻ cells and total YFP⁺ cells within these activated populations in spleen, lymph node and lung. Points represent individual mice. **E.** UMAP of the splenic CD4 T-cell scRNA-seq dataset coloured by originating CXCR5⁺YFP⁺, CXCR5⁻YFP⁺, PD-1⁺YFP⁻ or PD-1⁻YFP⁻ sorted population.

**Extended Data Figure 10.**
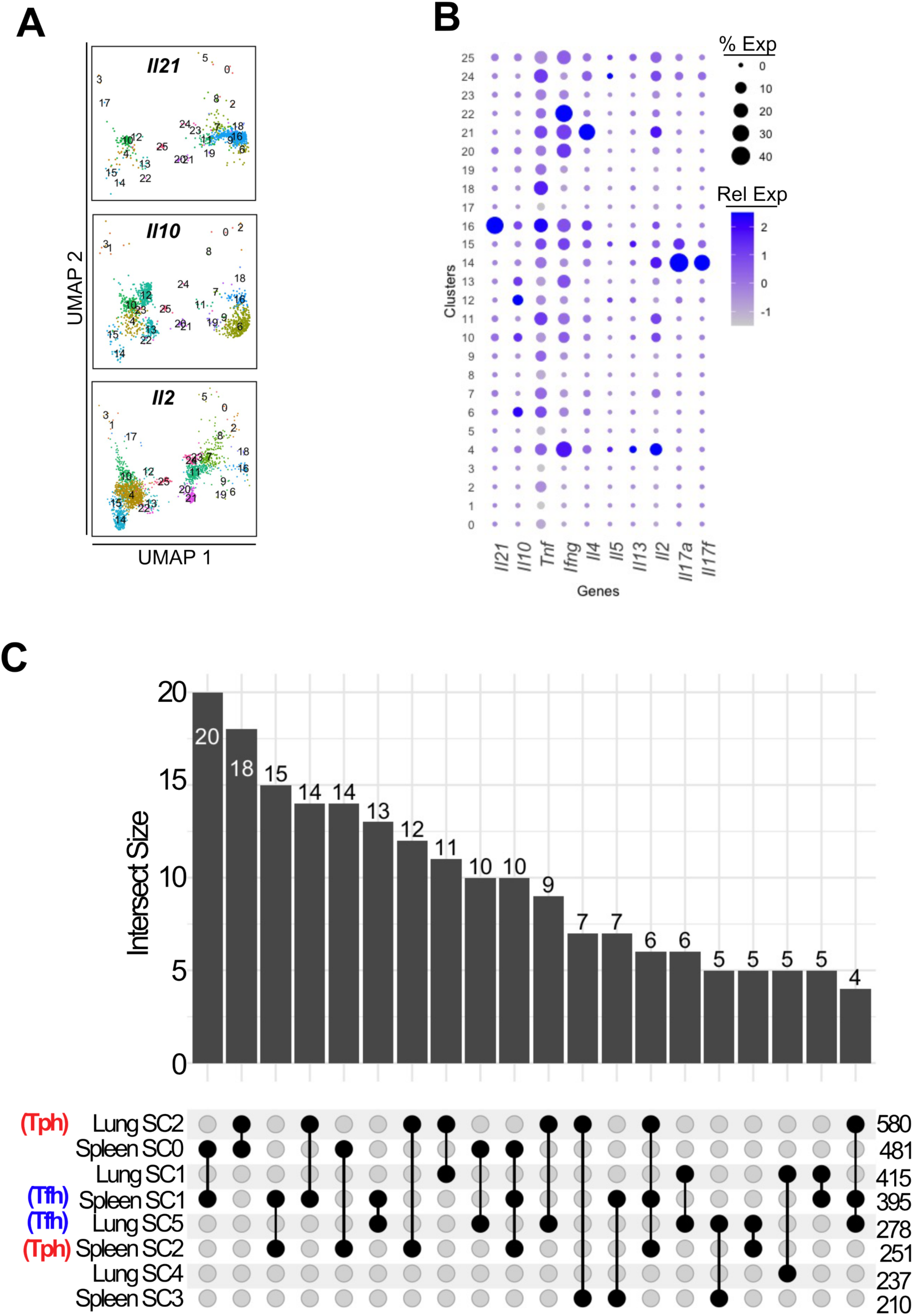
Cytokine expression across CD4 T cell clusters in CD4 T cells from Bcl6- sufficient or deficient mice. **A.** FeaturePlot expression of *Il21*, *Il10* and *Il2*. **B**. DotPlot of several different cytokines representative of different T cell functional states across all clusters, relative expression shown. C. Upset plot of TCR sharing between spleen cluster 16 subclusters and lung cluster 10 subclusters. Top 20 intersections are shown.

