## Supplementary Tables for "Clonal and developmental relationships between Tph and Tfh cells in patients with SLE and in murine lupus"

**Supplementary Table 1**

| CXCR5-PD1+ Spleen | CXCR5-PD1+ Lung | CXCR5+PD1+ Spleen | CXCR5+PD1+ Lung |
|---|---|---|---|
| Ccr8 | Gpr15 | Cd79b | Wdr90 |
| Nrn1 | Atp8b2 | Wdr90 | Apoe |
| Itgae | Fgl2 | Marf1 | Marf1 |
| Rpl7a | Cish | Apoe | Cd79b |
| Scin | Nrn1 | Padi4 | Cd74 |
| Foxp3 | Rpl7a | Cd74 | Psme2b |
| Gpr15 | Cxcr6 | Kyat3 | Dnase1l3 |
| Areg | Foxp3 | Smox | Mbd5 |
| Id2 | Lcmt2 | Cxcr5 | Zfp68 |
| Psmb9 | Sertad3 | Lpp | Cd22 |
| Ahr | Ptpn4 | Cd22 | Rpl35rt |
| Glt28d2 | Ccr8 | Pou2af1 | Cxcr5 |
| Klrd1 | Itgae | Egln2 | Irf8 |
| Penk | Il18r1 | Atp1a3 | Ecel1 |
| Ccrl2 | Nrf1 | Ms4a1 | S1pr2 |
| Serpinc1 | Il2ra | Stk19 | Arnt2 |
| Adam17 | Ahr | Cd59b | Stard5 |
| Spry1 | Tbx21 | Cd40 | Pou2af1 |
| Fgl2 | Ccl4 | Coro2b | Ccdc127 |
| Tnfrsf1b | Klrk1 | Abcc3 | Dock8 |
| Atp8b2 | Cd59b | Nebl | Padi4 |
| Slc28a2b | Vim | Psme2b | Dynlt1b |
| Thsd7a | Spry1 | Bcl6 | Zmpste24 |
| Ttbk1 | Havcr2 | Egln3 | Adarb2 |
| Ints6 | Plscr1 | Zfp68 | Ccdc62 |
| Zbtb32 | Il18rap | Swap70 | Fuca2 |
| Lamc1 | Adam17 | Ank2 | Dcp1b |
| Nfil3 | Ell2 | Myb | Atcay |
| Rbpj | Plagl1 | Spib | Stk40 |
| Axl | Nr3c1 | Irf8 | Pfn2 |
| Adamts6 | Impa2 | Slc26a11 | Bcl6 |
| Ptms | Irf4 | Dnase1l3 | Tspyl3 |
| Bace2 | Lmna | Zfp704 | Ttbk1 |
| Nudt4 | Ccr2 | Dennd4c | Egln3 |
| Ppp1r16a | Itgb1 | Tjp3 | Zfp944 |
| Ebi3 | Furin | Lmo4 | Bpgm |
| Trf | Uhrf2 | Fam43a | Fbxo33 |
| Tnfrsf9 | Fasl | Nav2 | Unc93b1 |
| Fasl | Cc2d1a | Lrif1 | Asb2 |
| Arrb2 | Serpina3g | Ikzf3 | Casp7 |
| Gata3 | Cldnd1 | Esm1 | Nup188 |
| Cd200 | Dusp2 | Arc | Elmo3 |
| Gabarapl1 | Plec | Dclre1c | Vrk2 |
| Coro2a | Tnfrsf9 | Prrt2 | Spp1 |
| Fhip1a | Sec24d | Zfr2 | Gnptab |
| Ptprs | Igsf8 | Ly6c2 | Ctsz |
| Bivm | Tiam1 | Itga7 | Cpq |
| Tnfsf10 | Knl1 | S1pr2 | Scd2 |

| | | | |
|---|---|---|---|
| Acsbg1 | Ptgr3 | Tspyl3 | Pfkfb3 |
| Niban1 | Decr1 | Zfp882 | Scpep1 |
| Rab2b | Tatdn1 | Nab1 | Tbc1d4 |
| Tmem106a | Smad3 | Sec14l1 | Gna12 |
| Cnrip1 | Tle3 | Smim3 | Sec14l1 |
| Klrg1 | Ctla2a | Fas | Zeb2 |
| Epcam | Csgalnact1 | Vrk2 | Il1r2 |
| Dstn | Gata3 | Ifi30 | Dpp8 |
| Cd83 | Fryl | Ttll12 | Mbnl3 |
| Pick1 | Runx2 | Itpr2 | Cisd1 |
| Tmem62 | Antxr2 | Fads6 | Dhrs1 |
| Tnfrsf4 | Alcam | Pla2g6 | Esm1 |
| Prkca | Lgals3 | Cdh23 | Nab1 |
| Irf4 | Serpinb9 | Nup188 | Ndufaf3 |
| Nr4a2 | Rpl3 | Ccr4 | Mcub |
| Cobll1 | Ebi3 | P2rx7 | Ddx55 |
| Slc52a3 | Ptpa | Hif1a | Galnt9 |
| Dtx4 | Nr3c2 | Zdhhc5 | P2rx7 |
| Gpr160 | Pros1 | Cdc25b | Tmem176b |
| Plscr1 | Axl | Srgap2 | Spaca6 |
| Maged1 | Ggct | Trbc1 | Rgs16 |
| Kdm5d | Serpinb6b | Rap1gap2 | Cyfip1 |
| Bmyc | Creb1 | Primpol | Cdh23 |
| Gpm6b | Cibar1 | Meis3 | Klc1 |
| Spp1 | Ccdc50 | Nek6 | Rarg |
| Slc16a3 | Tnfrsf18 | Tm6sf1 | Upf1 |
| Lratd1 | Zfyve26 | Asb2 | Acd |
| Wls | Tmem123 | Nudt22 | Golph3l |
| Bhlhe40 | Etv6 | Dck | Dap |
| Dok2 | Ogt | Txndc5 | Bmp7 |
| Tert | Cmtm3 | Chchd2 | Nfia |
| Phlpp1 | Ildr1 | Slc9a9 | Tmem176a |
| Pdcd1lg2 | Slfn5 | Lyn | Scamp2 |
| Synpo | Chsy1 | Scpep1 | Cd38 |
| Pmvk | Apex2 | Leng9 | Cdk2ap1 |
| Susd2 | Crtap | Ttc39c | Fas |
| Rpl36a | Nrp1 | F2rl1 | Med15 |
| Mdfic | Rab37 | Unc93b1 | Fkbp2 |
| Tmbim1 | Arrb2 | Lrrn3 | Fam43a |
| Sh3bgrl | Anxa6 | Sptb | Il10 |
| Cc2d1a | Plp2 | Dusp7 | Nfkbib |
| Ybx3 | Tnfrsf1b | Ehd1 | Vps26c |
| Nr3c2 | Atxn1 | Cyp4f16 | Nampt |
| Cacna1d | Nkg7 | Nemp2 | Rnf214 |
| Mecp2 | Tnfsf8 | Dennd2d | Slc22a15 |
| Nrp1 | Ehbp1l1 | Baiap3 | Vps13a |
| Gem | Maz | Tnfrsf8 | Stx11 |
| Alcam | Penk | Mical2 | Adamts6 |
| Ptger4 | Samsn1 | Gpr155 | Rab11a |
| Apbb1 | Lrrn3 | Ggt7 | Mfap1a |

| | | | |
|---|---|---|---|
| Nlrx1 | Cd200 | Fuca2 | Rnf169 |
| U2af1 | Creb3l2 | Trp53bp1 | Setbp1 |

**Supplementary Table 2. Human-derived Tph and Tfh signatures**

| Tph | Tfh |
|---|---|
| Lag3 | Gng4 |
| Slc27a2 | Cxcl13 |
| Havcr2 | Ascl2 |
| Cxcr6 | Ksr2 |
| Myo7a | Il21 |
| Rdh10 | Pvalb |
| Nfil3 | Cebpa |
| Ankrd35 | Cd200 |
| Vdr | Cxxc5 |
| Gna15 | Pdcd1 |
| Fam3c | Pou2af1 |
| Ccl4 | Fzd3 |
| Cxcl13 | Tox2 |
| Sdc4 | Cd79a |
| Pdcd1 | Fam43a |
| Camk1 | Xxylt1 |
| Tnfrsf18 | Smad1 |
| Furin | 5330417C22Rik |
| Ctla4 | Bcl6 |
| Otud1 | Ica1 |
| Slc16a3 | Scgb3a1 |
| Arl3 | Cxcr5 |
| Gabpb1 | Nfatc1 |
| Prdm1 | Igfbp4 |
| Dusp4 | Gfod1 |
| Znrf1 | Ptpn11 |
| Snx9 | Cep128 |
| Ankrd28 | Tigit |
| Gde1 | Id3 |
| Runx2 | Tox |
| Sytl2 | Itm2a |
| Hopx | Sardh |
| Nudt16 | Cd59b |
| F2r | Btla |
| Tbxas1 | Ier5l |
| H2-Eb1 | Coro1b |
| Tnfrsf4 | |
| Rbpj | |
| Lgals1 | |

**Supplementary Table 3. Human C5 to mouse signatures**

Tox
St8sia1
Ttn
Sesn3
Fut8
Tigit
Cxcr3
Ldlrad4
Slamf6
Gtdc1
Rgs1
Itgb1
Itm2a
Smc4
Pam
Tbc1d4
Jak2
Ubash3b
Dgkh
Ibtk
Gypc
Tspan14
Dip2b
Sell
Wdfy1
Fam117b
Coro1b
Tbl1xr1
Atp8a1
Tmem245
Akap11
Gbp3
Ogt
Ucp2
Fkbp5
Gm5601
Zfp831
Slc9a9
Mis18bp1
Ptprj
Icos
Cnn2
Gbp2
Ipcef1
Cotl1
Nin
Nr3c1
Elmo1

**Supplementary Table 4. scRNA-seq-derived BCH mouse signatures**

| Spleen BCH | Lung BCH |
|---|---|
| Stfa3 | Gm34084 |
| 1810009J06Rik | Kcnc2 |
| Gm2663 | Ccser1 |
| Spp1 | Pdzrn3 |
| Edn3 | Wnt3 |
| Rph3a | Adarb2 |
| Il21 | Gpm6b |
| Vdr | Flrt2 |
| Gm15482 | Gm14718 |
| S100a5 | Chn1 |
| Pdzrn3 | Edn3 |
| Chn1 | Mmp10 |
| Tnc | Cd109 |
| Ehd2 | Plagl1 |
| 8030451A03Rik | Sostdc1 |
| Ascl2 | Bace2 |
| Kcnc2 | 8030451A03Rik |
| Fbn2 | Gna14 |
| Lpar3 | Gm43057 |
| Myh10 | Fam43a |
| Padi4 | Ypel4 |
| Flrt2 | Csgalnact1 |
| Pcsk1 | Gm26686 |
| Akr1b8 | Crppa |
| Sdk1 | Sh3rf2 |
| Rgs8 | Ralyl |
| Ifi204 | Angptl2 |
| Angptl2 | Twist2 |
| Ralyl | Vdr |
| Igfbp7 | Sptb |
| Fam20a | Padi4 |
| Gm12534 | Cebpa |
| Gm43057 | Csta2 |
| Gm14718 | Ascl2 |
| Exph5 | Gm2663 |
| Stc2 | Synpo |
| Zbtb32 | Pcsk1 |
| Sostdc1 | Ifi204 |
| Ccser1 | Cc2d2a |
| Cd109 | Ica1l |
| Tnfsf8 | Bcam |
| Sh3rf2 | Sh3gl3 |
| Cbfa2t3 | Fbn2 |
| Tspan2 | Il21 |
| Adipoq | Cybb |
| Kif15 | 2610204G07Rik |
| Gm31135 | Dmwd |
| Efnb2 | Shtn1 |

| | |
|---|---|
| Ica1l | Pkn3 |
| Plagl1 | Cd200 |

**Supplementary Table 5. scRNA-seq-derived Tfh Tph mouse signatures**

| Spleen Tph signatures | Spleen Tfh signatures |
|---|---|
| Nrn1 | Tspan5 |
| Bace2 | Evl |
| S100a4 | Ighm |
| Irf4 | Trim59 |
| 1700019D03Rik | 2010016I18Rik |
| Bhlhe40 | Bzw2 |
| Ahnak | Sh2d1a |
| Anxa6 | Pde7a |
| Alcam | Dap |
| Cd83 | Cited2 |
| Nfkbia | Znrf1 |
| Tnfrsf1b | Lef1 |
| Nudt4 | Abhd8 |
| Plec | Id3 |
| Id2 | Prmt3 |
| Cd200 | 4930417O13Rik |
| Capg | Ephx1 |
| Tnfrsf4 | Pdcd1 |
| Esyt1 | Camk4 |
| Rgs1 | Rnaset2a |
| Rbpj | Slc9a9 |
| Vim | Thada |
| Plscr1 | Ces2c |
| Glrx | Dennd2d |
| Spry1 | Cdc25b |
| Casp3 | Ubac2 |
| Prnp | Asb2 |
| S100a6 | Trerf1 |
| Slamf1 | P2rx7 |
| Sms | Rgs10 |
| Podnl1 | Tox2 |
| Nfkb2 | Pou2af1 |
| Ccnd2 | Cxcr5 |
| Klf2 | Nav2 |
| Sdf4 | Cebpa |
| Egr1 | Igkc |
| Ier2 | Ppp1r14b |
| Il7r | Fam43a |
| Lag3 | Dck |
| Capn2 | Matk |
| Prelid2 | Bcl6 |
| Btg3 | Art2b |
| Serpina3g | Rflnb |
| Pdcd1lg2 | Ascl2 |
| Bst2 | Nsg2 |
| Atxn1 | Pde2a |
| Nfil3 | Tubb2a |
| Mndal | Cxxc5 |

| | |
|---|---|
| Ptms | Atp1b1 |
| Crip1 | Gm30211 |
